# Cell-Fate Determination from Embryo to Cancer Development: Genomic Mechanism Elucidated

**DOI:** 10.1101/637033

**Authors:** Masa Tsuchiya, Alessandro Giuliani, Kenichi Yoshikawa

## Abstract

The elucidation of the how and when of a cell-fate change asks for a physically reasonable mechanism allowing to achieve a coordinated switching of thousands of genes within a small and highly packed cell nucleus. We previously demonstrated that whole genome expression is dynamically self-organized through the emergence of a critical point. Furthermore, it has been confirmed that this happens at both the cell-population and single-cell level through the physical principle of self-organized criticality.

In this paper, we further examine the genomic mechanism which determines cell-fate changes from embryo to cancer development. The state of the critical point, acting as the organizing center of cell-fate, determines whether the genome resides in a super- or sub-critical state. In the super-critical state, a specific stochastic perturbation can spread over the entire system through the ‘genome engine’ - an autonomous critical-control genomic system, whereas in the sub-critical state, the perturbation remains at a local level. We provide a consistent framework to develop a biological regulation transition theory demonstrating the cell-fate change.

## I. Introduction

A mature mammalian somatic cell can reprogram its state and consequently acquire a very different gene expression profile through few reprogramming stimuli [Takahashi, K., Yamanaka, S., 2016]. Such drastic state change involves the coherent on/off switching of thousands of functionally heterogeneous genes [MacArthur, B. D., et al., 2009]. However, there are fundamental physical difficulties in achieving such large-scale coordinated control on a gene-by-gene basis. These difficulties become more evident in a situation where 1) there is a lack of a sufficient number of molecules to reach a stable thermodynamic state (i.e., breakdown of the central limit theorem) and 2) a consequent stochastic noise due to the low copy numbers of specific gene mRNAs, inducing a substantial instability of genetic product concentrations. [Raser, J. M., O’Shea, E. K., 2005; Yoshikawa, K., 2002].

In our previous studies [Tsuchiya, M., et al., 2014-2017; Giuliani, A., et al., 2018, Zimatore, G. et al, 2019], we demonstrated the self-organization of the whole genome expression constitutes a ‘physically motivated’ alternative to the gene-specific regulation at both single cell and cell population scales. The mechanism of self-organization, through global genome reprogramming, eliminates the need of physically unfeasible gene-by-gene expression control.

The core of the self-organization mechanism is the presence of massive system changes elicited by ‘apparently minor’ external causes. To address this problem, Per Bak and colleagues [Bak, P., et al., 1987] proposed self-organized criticality (SOC; the Bak-Tang-Wiesenfeld sandpile model). SOC is a general theory of complexity that describes self-organization and emergent order in non-equilibrium systems (thermodynamically open systems). Self-organization, often with the generation of exotic patterns, occurs at the edge between order and chaos [Langton, C. G., 1990; Kauffman, S. A., 1993]. Further description of SOC is included in [Jensen, H. J. 1998; Marković, D., Gros, C., 2014] and for a review of the literature on criticality, see [Muñoz, M. A., 2018].

SOC builds upon the fact that the stochastic perturbations initially propagate locally (i.e., sub-critical state); however, due to the particularity of the disturbance, the perturbation can spread over the entire system in a highly cooperative manner (i.e., the super-critical state). As the system approaches its critical point, global behavior emerges in a self-organized manner. The coordinated character (and possible self-organization) of the process stems from the so called ‘domino effect’ present in all the biological signaling (e.g., in allosteric effect, see: [Wagner, J. R., et al., 2016])) where microscopic local effects generalize to the entire system spreading along ‘preferential pathways’.

The above-depicted classical concept of SOC explained above has been extended to propose a conceptual model of the cell-fate decision (critical-like self-organization or rapid SOC) through the extension of minimalistic models of cellular behavior [Halley, J. D., et al., 2009]. The cell-fate decision-making model considers gene regulatory networks to adopt an exploratory process, where diverse cell-fate options are generated by the priming of various transcriptional programs. As a result, a cell-fate gene module is selectively amplified as the network system approaches a critical state. Such amplification corresponds to the emergence of long-range activation/deactivation of genes across the entire genome.

We adapted SOC paradigm at cell-fate decision and investigated whole genome expression and its dynamics to address the following fundamental questions:

- Is there any underlying principle that self-regulates the time evolution of whole-genome expression?
- Can we identify a peculiar genome region guiding the super-critical genome and determining cell fate change?
- Can we rely on a universal mechanism to grasp the how and when of cell-fate change occurs?

Our previous studies of self-organization with critical behavior (criticality) ([Tsuchiya, M., et al., 2016]) differ distinctly from the classical and extended SOC models in regard to the following issues:

1. **Self-organization**: When gene expression is sorted and grouped according to temporal variance of expression (normalized root mean square fluctuation: *nrmsf*; **Methods**), a transitional behavior of expression profile occurs in the ensemble of genes to exhibit coexistence of local critical states. These states include: 1) super-critical state, high temporal-variance expression; 2) near-critical state, intermediate variance expression; 3) sub-critical state, low variance expression, in which *nrmsf* acts as an order parameter of self-organization.
2. **Critical behaviors**: Two distinct critical behaviors emerged when gene expression values are sorted and grouped: 1) sandpile-type criticality and 2) scaling-divergent behavior. Sandpile criticality, where the sandpile-summit corresponds to a critical point (CP), is evident in terms of grouping according to expression fold-change between two different time points, whereas scaling-divergent behavior emerges according to grouping by *nrmsf* (refer to Fig.1 in [Giuliani, A., et al., 2018]). Criticality allows for a perturbation of the self-organization due to change in signaling by external or internal stimuli into a cell to induce a global impact on the entire genome expression system. The scaling-divergent behavior may reveal a quantitative relationship between the aggregation state of chromatin through *nrmsf* and the average expression of an ensemble of genes. The degree of *nrmsf* should be related to the physical plasticity of genomic DNA and the high-order chromatin structure.
3. **Material basis of SOC control**: Chromatin remodeling is the material basis of the SOC control of genome expression. Additionally, a cell-fate guided critical transition occurs on genome expression through coordinated change of local chromatin folding [Zimatore, G. et al, 2019].
4. **Genome-engine**: A highly coherent behavior of low-variance genes of the sub-critical state generates a dominant cyclic expression flux with high-variance genes of the super-critical state through the cell environment to develop autonomous critical control system. The local sub-critical state as the generator of autonomous SOC control guides the cell-fate change versus the non-autonomous classical SOC model [Halley, J. D., et al. 2009].
5. **Cell-fate change**: Cell-fate change occurs through erasure of the initial-state sandpile critical behavior (criticality). SOC control of overall gene expression (i.e., initial-state global gene expression regulation mechanism) is eliminated through the erasure of an initial-state criticality. This suggests that the critical gene ensemble of the CP plays a significant role in determining the cell-fate change.

In this report, we update our previous findings by elucidating the specific critical genome region (i.e., critical point: CP) for cell-fate control involved in its critical transition, and examine a potential genomic mechanism underlying cell-fate change. Our report is organized as described in **Figure 1**:

**Figure 1:**
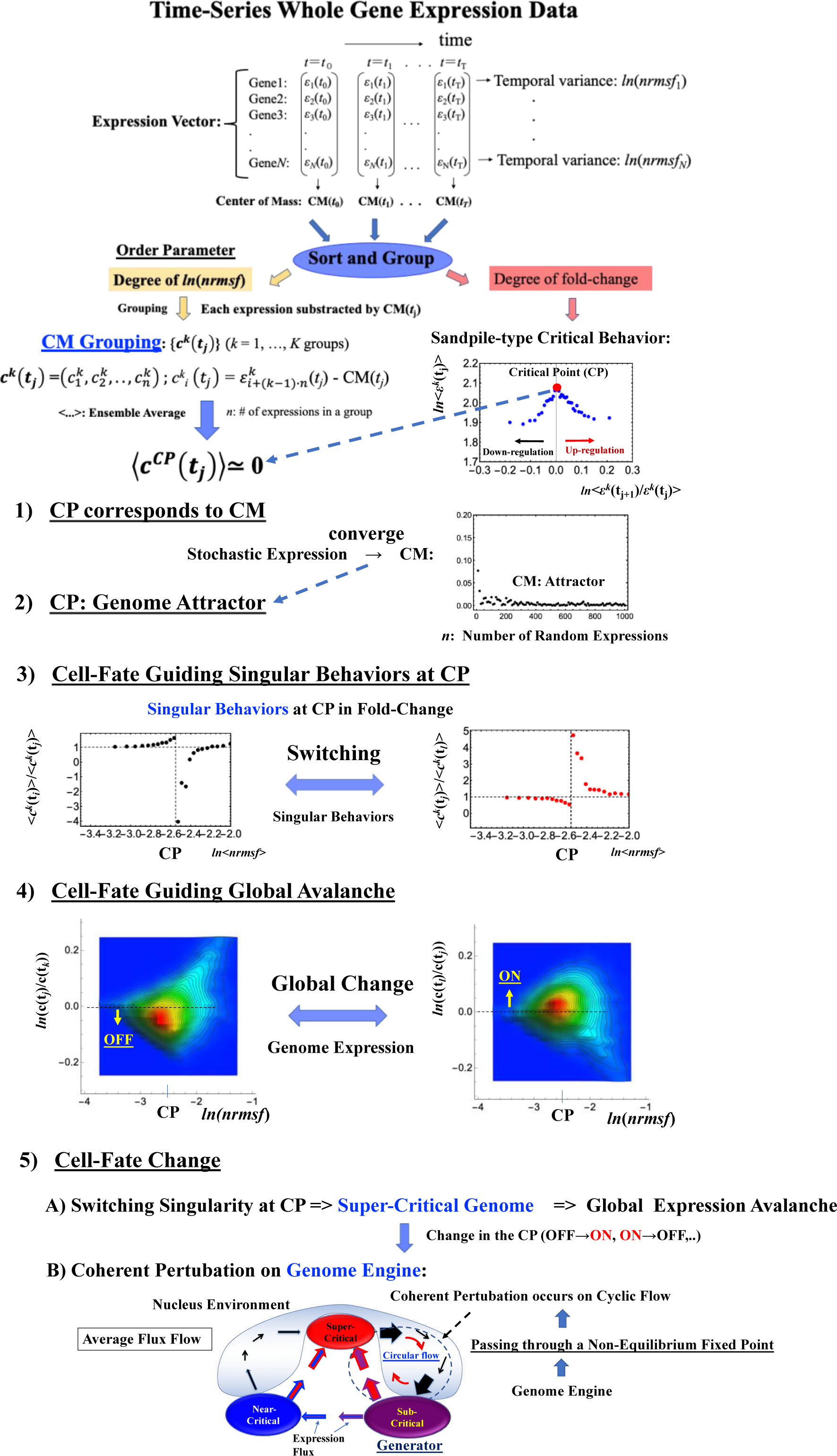
Schematic Representation of Genomic Mechanism for Cell-Fate Change. **1-4)** The CP corresponds to the center of mass (CM) and represents a specific set of critical genes acting as genome-attractor. The CP has an activated (ON) or deactivated (OFF) state. The ON/OFF switch of the CP state occurs through switching its singular behaviors, change in critical transition of specific critical gene set. In OFF state of the CP, the stochastic perturbations propagate locally, but when the particularity of the disturbance activates the CP, i.e., the genome becomes super-critical, the perturbation can spread over the entire system in a highly cooperative manner [Tsuchiya, M, et al., 2019]. The role of chromatin remodeling plays as the material basis of the SOC control of genome expression [Zimatore, G. et al, 2019]. **5)** Due to the CP acting as the genome-attractor, self-organization of gene expression develops an autonomous critical-control genomic system (genome-engine) through the formation of dominant cyclic flux between local critical states (distinct expression domains according to the degree of *nrmsf*), where the local sub-critical state is the generator (see details in [Tsuchiya, M, et al., 2016]). Coherent perturbation on the genome engine through the change in the CP (ON to OFF, OFF to ON, etc.) drives cell-fate change. Before the cell-fate change the genome (expression) system passes through a non-equilibrium fixed point (stable point of thermodynamically open system). These five points support the development of a biological regulation transition theory.

1) The CP corresponds to the center of mass (CM(*t*_j_): average value of whole expression at *t* = *t*_j_) (**section IIA**). 2) The dynamics of the center of mass of any stochastic expression converges to that of the whole expression (i.e., (CM(*t*_j_)). This shows that the CP is the genome-attractor (**section IIB**). 3) The switching singular behaviors of the CP transforms the genome into a super-critical state (i.e., super-critical genome). This in turn induces a ‘global expression avalanche’, which is revealed by 4) the probability density profile (PDF) of the whole expression. 5) Lastly, cell-fate change occurs after the genome becomes a ‘super-critical genome’ (**sections IIC-E**) and after the genome passes over a stable point (non-equilibrium fixed point) of the thermodynamically open system (**section IIG**). This passing indicates symmetry breaking inducing coherent perturbation on the genome-engine (**section IIF**). These five points constitute a framework for developing a biological regulation transition theory.

## II. Results

### A. Fixed Critical Point (CP): A Specific Group of Genes Corresponding to the Center of Mass of Whole Genome Expression

The existence of a critical point (CP) is essential in determining distinct response domains (critical states) [Tsuchiya, M., et al., 2016]. To generate a unified model of biological regulation, we must go in depth into specific features of the CP in terms of ‘*sandpile criticalit*y’ (**Figures 2A**). Sandpile criticality emerges when the whole genome expression is sorted and grouped according to fold-change in expression between two different time points (e.g., between *t* = 0 and *t* =10min). For the same groupings, *nrmsf* value of the CP in HRG-stimulated MCF-7 cancer cells (population level) is estimated (*ln*<*nrmsf*> ∼ -2.5: **Figure 2B**).

**Figure 2:**
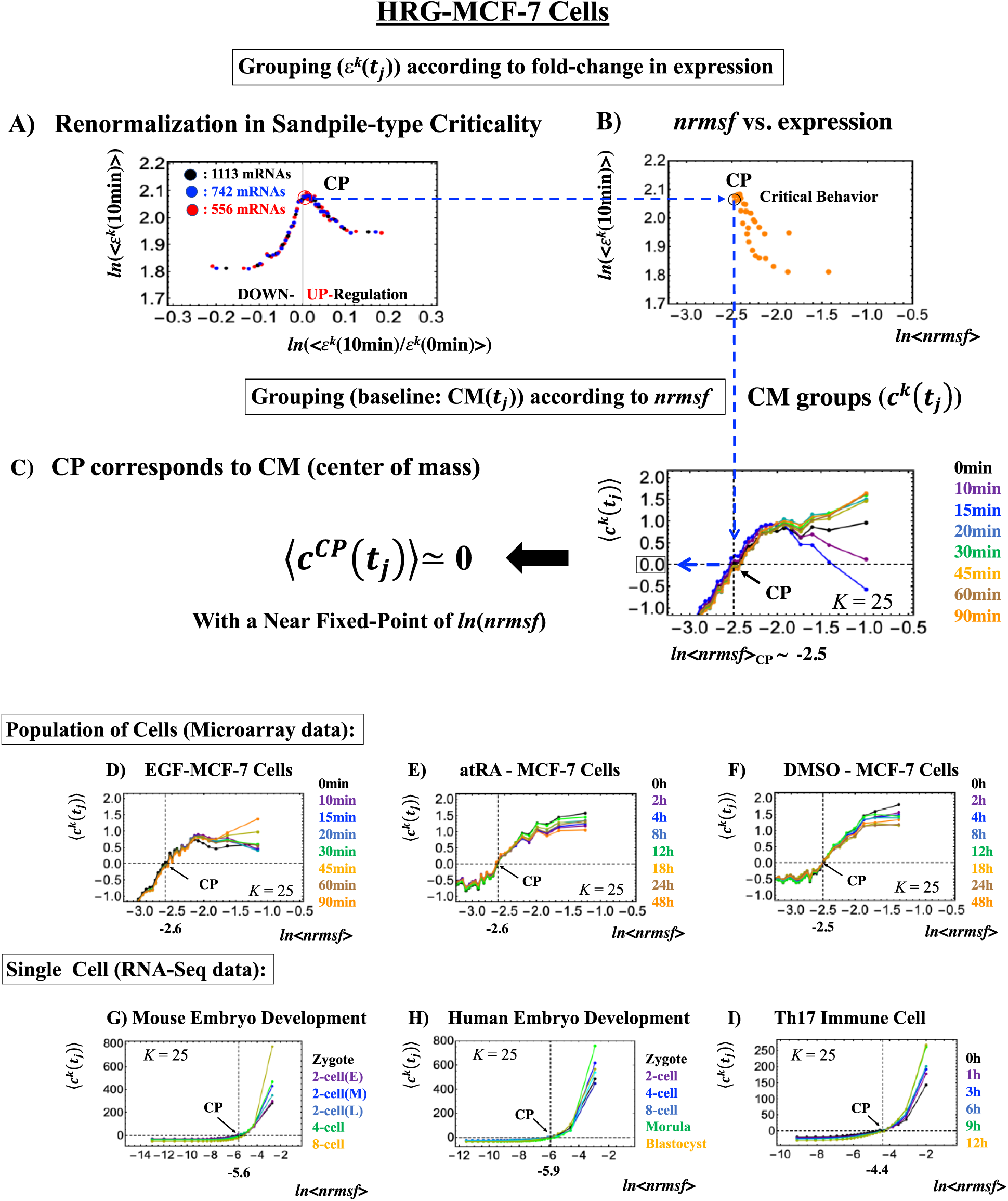
The fixed CP corresponding to the center of mass (CM) of genome expression: **A)** Different number of groups (*ε*^*k*^(*t*), *k* = 1,2…,*K*: non-CM grouping; black: *K*= 20; blue: 30; red: 40 groups; each group contains 1113, 742, and 556 mRNA species respectively) in HRG stimulated MCF-7 cells follow the same sandpile-type critical behavior. This reveals the existence of scaling behaviors (i.e., renormalization). **B)** *ln*<*nrmsf*> value of the CP: logarithm plot of average *nrmsf* value of group, *ln*<*nrmsf* > vs. average expression value, *ln*⟨*ε*^*k*^(10*min*)⟩ shows that *ln*<*nrmsf*>_CP_∼ -2.5 in sandpile type criticality, where *ε*_*k*_(10min) represents expression value of the *k*^th^ group (ordered from high to low *nrmsf* value) at *t* = 10min and <…> represents ensemble average of group expression. Grouping for **A)** and **B)** is ordered according to fold-change in expression at 0-10min. **C)** Grouping (*c*^*k*^(*t*)) (CM grouping) of whole expression (baseline: CM(*t*_*j*_)) according to *nrmsf* reveals that this can be considered a fixed point and corresponds to the CM of genome expression (whole expression). This fact is true for both the population (**C-F**) and single-cell levels (**G-I**). *K* represents the number of groups with *n* number of elements (**C, D**: *n* = 891 mRNAs; **E, F**: *n* = 505; **G**: *n* = 685 RNAs; **H**: *n* = 666; **I**: *K* = 25, *n* = 525; coloring: **Methods**).

Our study on HRG-stimulated MCF-7 cancer cells demonstrated that the temporal group correlation (between-groups correlation) along the order parameter (*nrmsf*) reveals a focal point (FP) when we consider the center of mass (CM) of whole expression (changing in time) as a reference expression point (see Fig. 5B in [Tsuchiya, M., et al., 2015]). The grouping (baseline as the CM(*t*_j_)) according to the degree of *nrmsf* is called ***CM grouping***, *c*^*k*^(*t*_*j*_) (*k*^th^ group; *k* =1,.., *K*), where grouping from the CM distinguishes from that of non-reference, *ε*^*k*^(*t*).

**Figure 3:**
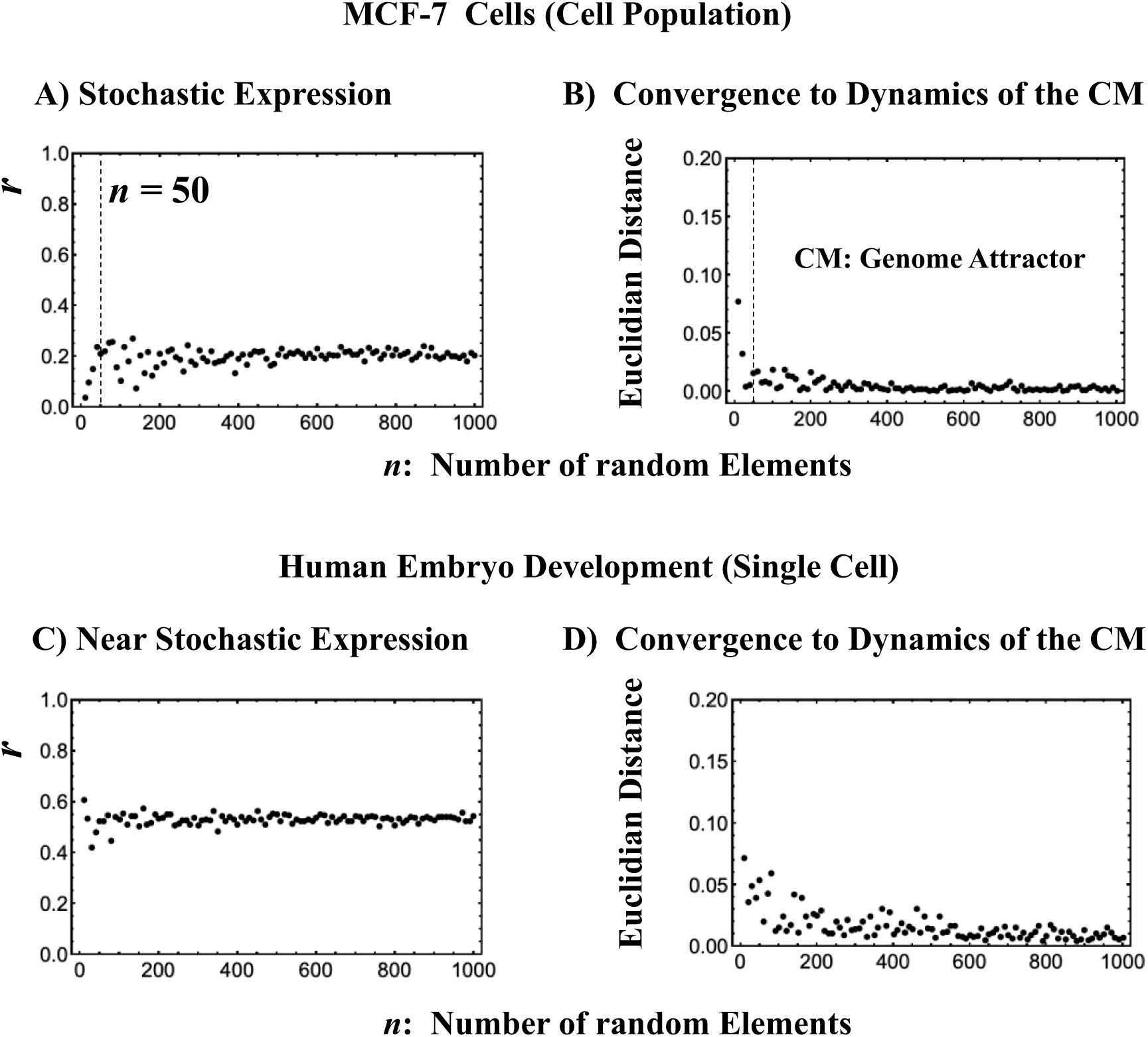
The center of mass (CM) acting as genome-attractor. MCF-7 cells (Microarray data: **A** and **B**) and Human embryo development (RNA-Seq data: **C** and **D**). The *x*-axis represents the number of randomly selected single gene expression values in all the four panels while y-axis reports the average Pearson correlation. Panels **A)** and **C)** illustrate that bootstrapping highlights a low pairwise Pearson correlation in randomly selected groups of expression (computed over experimental time points with 200 repetitions). This confirms the stochastic character of gene expression. The removal of all genes with zero expression value at a time point does not change their correlation behavior (**C)** (similar behavior is also observed in mouse embryo case; data not shown). The higher correlation makes the convergence slower than in the cancer case (**B**). Panels **B)** and **D)** illustrate the convergence of the CM of a randomly selected group of expression to the CM of whole expression over experimental time points with 200 repetitions as the number of selected genes increases (*x*-axis). The CP, the CM of the whole expression according to *nrmsf* acts as the genome-attractor. Note: as for the rest of biological regulations throughout this report, similar behaviors are observed. The dashed lines indicate in in graphs **A** and **B** that coherent behaviors emerge at *n* = 50 single gene expression values.

**Figure 4:**
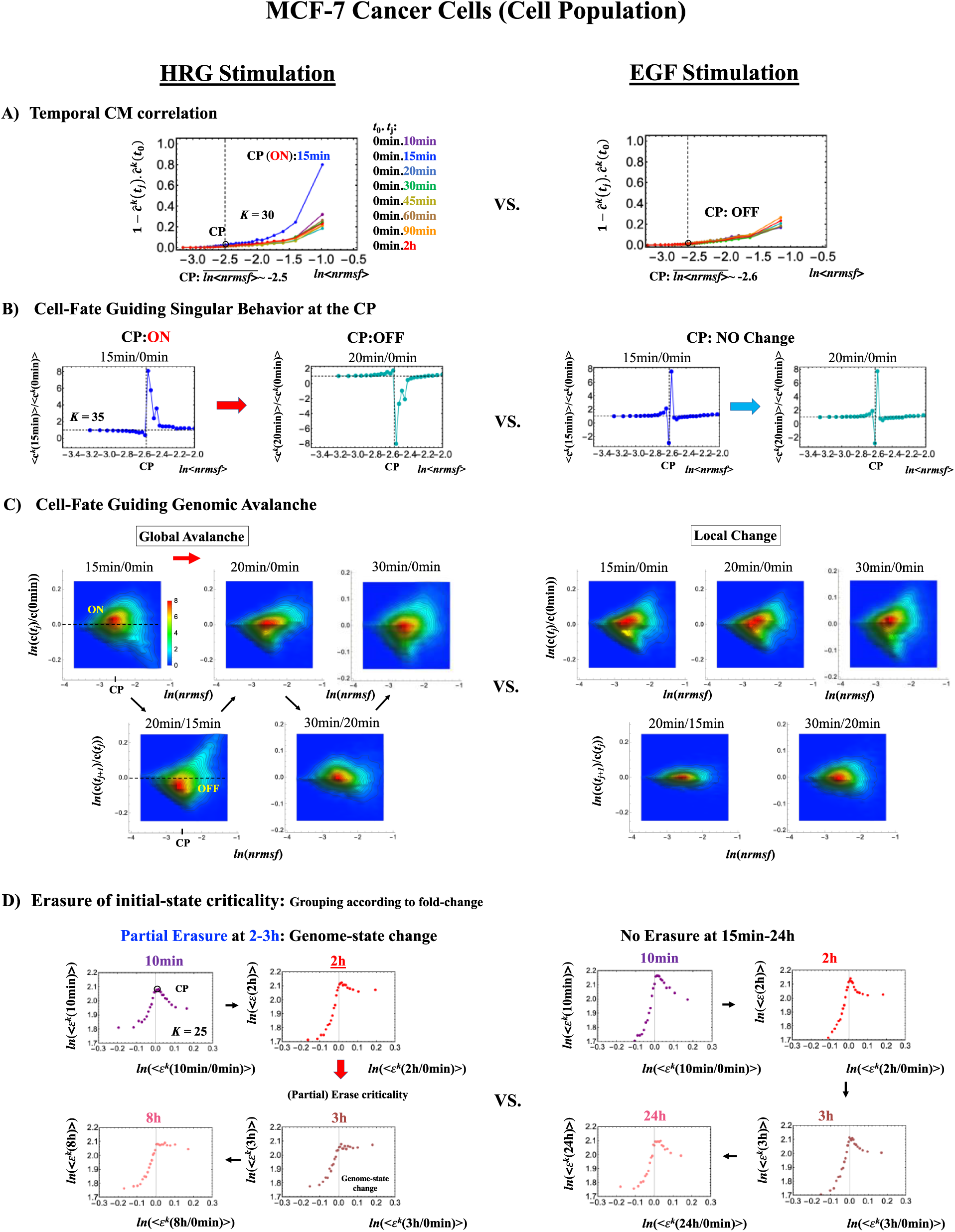
HRG- (left) and EGF-stimulated (right) MCF-7 cell-population responses. The CM correlation analyses (**Methods**) reported in this figure are relative to gene groups ordered along their relative *nrmsf* (*x-*axis): **A)** temporal CM correlation (**Methods**): 1 − ***ĉ***^*k*^(*t*_*j*_). ***ĉ***^*k*^(*t*_0_) (*y*-axis) is evaluated. At 15 min, the correlation response for HRG-stimulation is diverging from other responses, indicating that the CP is activated between 10 and 15 min for HRG. Whereas for EGF-stimulation, such distinct diverging correlation response does not occur (i.e., the CP is inactivated). Activation or inactivation of the CP marks the occurrence of cell-differentiation for HRG and only proliferation (no differentiation) for EGF. Panels **B)** and **C)** provide direct evidence of the ON/OFF state of the CP: **B)** The transition of the higher-order structure of the CP occurs for 15-20 min as shown in fold change from initial time point. Swelled coil state (dominant positive fold-change: *y*-axis) at 15min and compact globule (dominant negative fold-change) state at 20min respectively correspond to ON and OFF states of the CP (*K* = 30 groups with each group containing *n* = 742 mRNAs). Throughout this report, to make double in the number of groupings, a new group is added in the plot, where the group is created by combing two halves from the next neighbors of CM groups, and its average expression is evaluated. **C)** The probability density function (PDF; see **Methods**) of the whole expression (*z*-axis) shows that global expression avalanche occurs at 15-20min (coinciding with the change in the CP), where maximum probability density occurs around the CP with a positive value of natural log of fold-change (i.e., the CP is ON) at 15 min, whereas it becomes negative (OFF) at 15-20 min period. **D)** The (partial) erasure of the initial-state criticality (grouping according to fold-change in non-reference expression, *ε*^*k*^(*t*): *K* = 25: *n* = 891 mRNAs) occurs at 2-3hr for HRG (indication of genome-state change) and no erasure for EGF.

**Figure 5:**
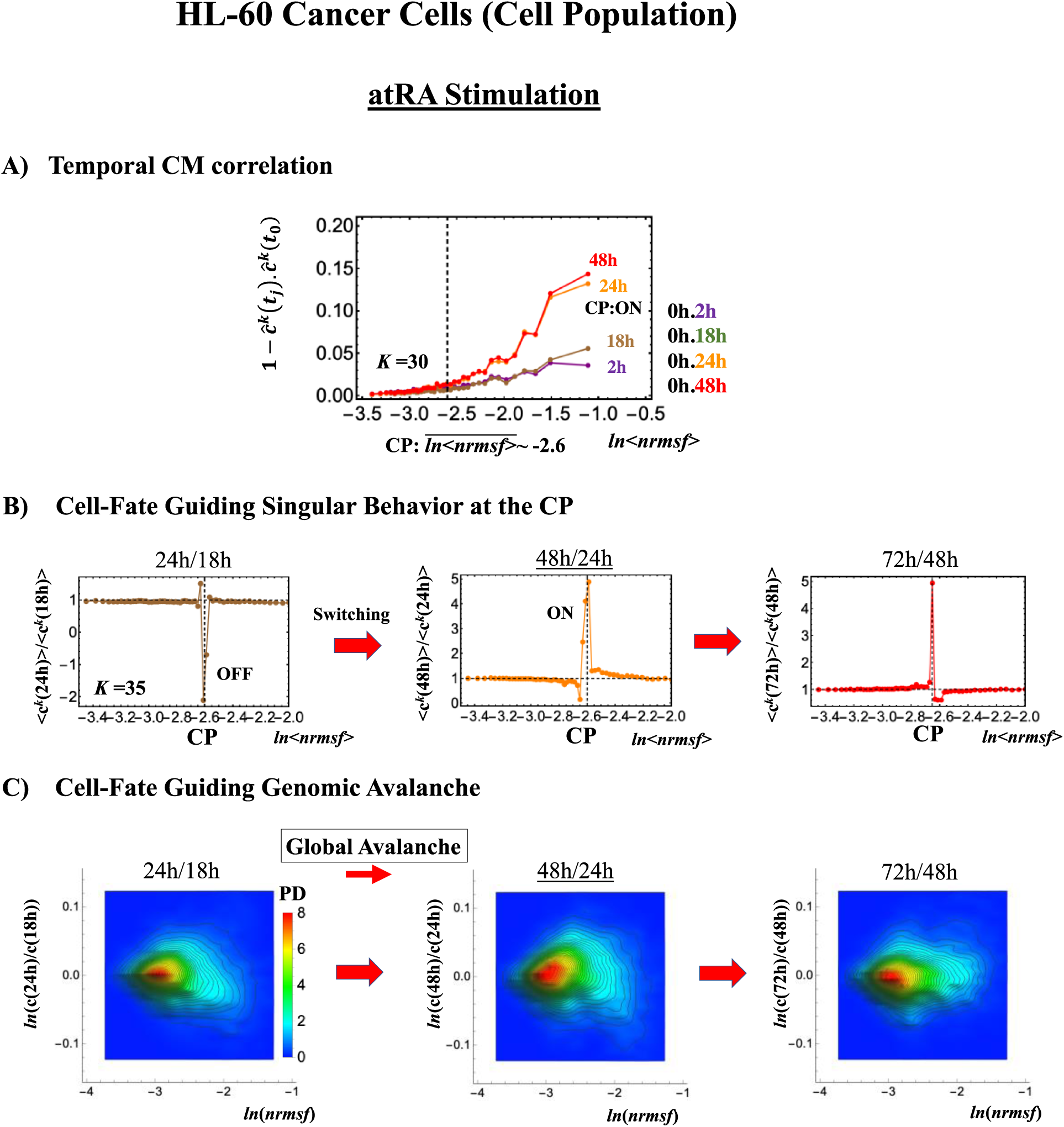
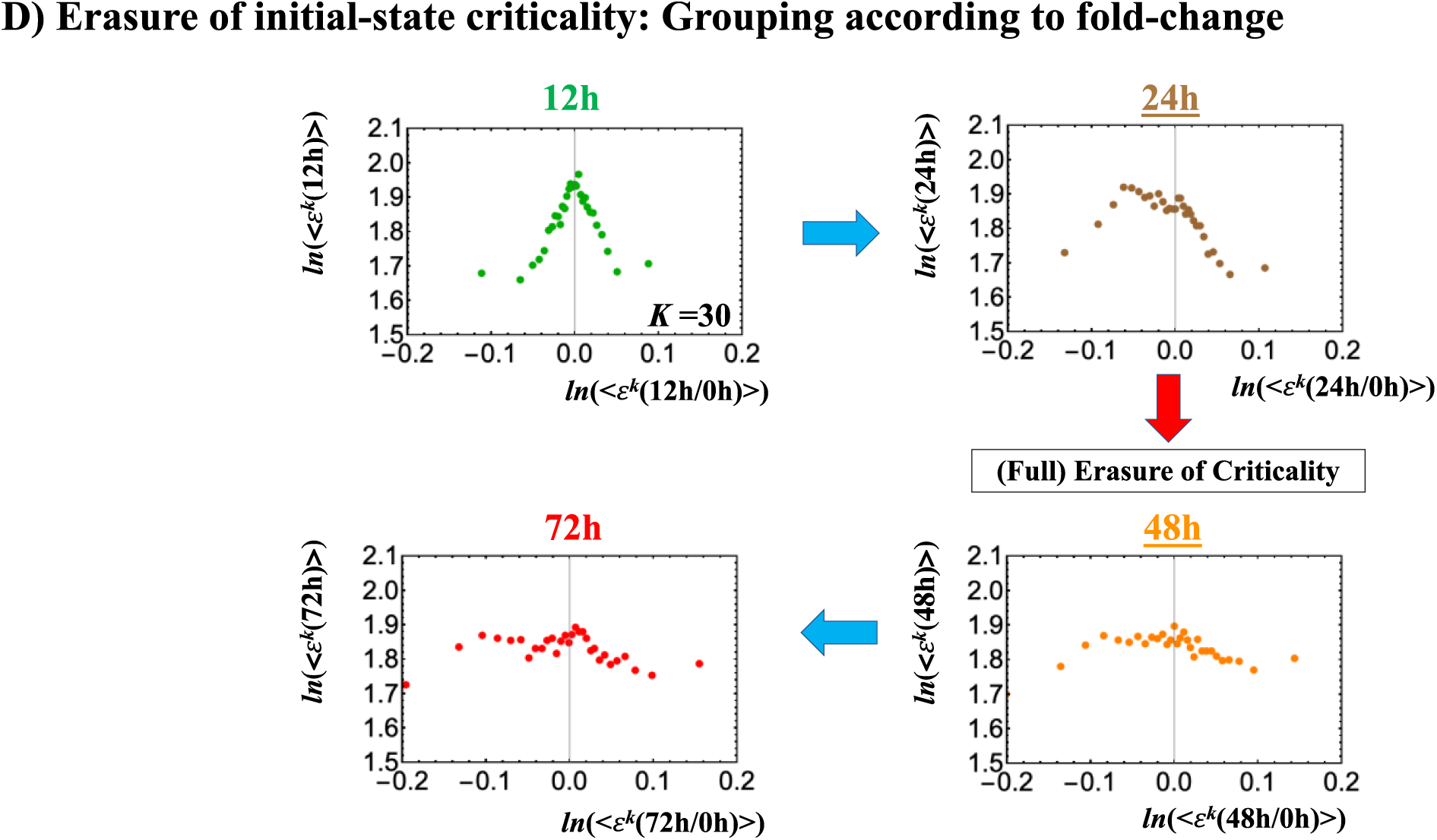
atRA-stimulated (right) HL-60 cell population. **A)** Activation of the CP occurs (*K* = 30: *n* = 420 mRNAs) at 24-48h for atRA shown by the temporal CM correlation response. **B)** Transition of the higher-order structure at the CP (*K* = 35: *n* = 360 mRNAs) is suggested to occur at 24-48h, when the initial-state criticality is erased (**D**). The switching of singular behaviors at the CP occurs with bimodal behaviors of fold-change around the CP. The dominant negative fold-change indicates OFF state (compact globule DNA) occurs at 18-24h, while the dominant positive fold-change (swelled coil DNA) occurs at 24-48h. **C)** Temporal change in PDF shows that global avalanche occurs from 18-24h (PDF: contracted) to 24-48h (swelled out) to 48-74h (contracted), which supports the switching behaviors of the CP (**B**). **D)** The full erasure of the initial-state criticality occurs at 24-48h, which indicates cell-fate change. Underlined numbers represent the timing for the cell-fate change.

Notably, as shown in **Figure 2C**, the CP corresponds to the zero-expression point in the CM grouping, which explains why the CP is a specific set of genes corresponding to the CM(*t*_*j*_) and the CP exists as an almost fixed point in terms of the order parameter, *nrmsf*. This feature holds for both single-cell and population data (**Figures 2 D-I**: refer to natural log of *nrmsf* value for different biological regulations). Therefore, we develop the correlation metrics based on the CM grouping (called **CM correlation**: **Methods**) to grasp how the whole expression can be self-organized through critical/singular behavior of the CP.

In [Zimatore, G., et al., 2019], we demonstrated that CP genes expression correspond to the attractor specific (i.e. the values typical of the specific cell kind analysed) values with almost null influence of fluctuation modes, consistently to their zero-expression point in CM grouping.

### B. Critical Point (CP) Acting as the Genome-Attractor: Mechanism for Genome-Wide Avalanche

The genome expression possesses coherent-stochastic behaviors (CSB) in which coherent behavior emerges from an ensemble of stochastic expression [Tsuchiya, M., et al., 2014-2017]. In CSB mode, gene expression is inherently stochastic but its dynamics follows the center of mass (CM) of expression. Interestingly, CSB is evident at both whole expression (**Figure 3**) and at specific critical states levels (Figs. 3B, D in [Tsuchiya, M., et al., 2017]).

To give a proof-of-concept of CSB, we performed a bootstrap simulation approach to catch two basic signatures of CSB:

1. Stochastic behavior of gene expression shows (relatively low) correlation convergence for randomly selected gene ensembles as the number of elements (*n*) is increased (**Figure 3A**),
2. The CM of randomly selected gene ensembles (with hundreds of repetitions for the gene ensembles) must dynamically converge to that of whole expression as the number of elements (*n*) is increased (**Figure 3B**).

This happens at both population and single-cell level (**Figures 3 C,D**). The existence of a threshold *n* at around 50 randomly picked genes [Censi, F., et al., 2011; Tsuchiya, M., et al., 2015] allows to reproduce CSB with a random choice of *N* genes with *N* > *n*, which is a further proof of the reliability of coherent-stochastic behaviors. The constancy of the minimum number of randomly selected genes to reach the convergence (**Figure 3**) is remarkable and suggests the presence of a sort of ‘percolation threshold’ reminiscent of the size of genetic networks operating in the system.

The convergence clearly reveals that the dynamics of the CM of genome expression describes an attractor of the dynamics of stochastic expression. Therefore, as shown in **Figure 3**, the CP, the CM according to the degree of *nrmsf*, acts as the *genome attractor*. Therefore, a change in the CP provokes a genome-wide avalanche over the entire genome expression. Whole expression follows the change in the CP - the origin of the coherent gene expression behaviors [Tsuchiya, M., e al., 2007].

Here, it is important to note that CSB does not stem from the law of large numbers under an equilibrium condition, rather than from an emergent property of open (non-equilibrium) system of the genome; e.g., a mixed critical state breaks the convergence to the CM, while each critical state exhibits CSB (Figs. 3D in [Tsuchiya, M., et al., 2017]).

### C. ON/OFF State of the CP: Cell-Fate Guiding and Global Avalanches

We consider cell-fate change equivalent to a genome-state change (refer to biological discussions in [Tsuchiya, M., et al, 2016]). The genome-state change occurs in such a way that the initial-state SOC control of whole genome expression, which is in charge of keeping largely invariant global gene expression profile, is destroyed through the erasure of an initial-state sandpile criticality [Tsuchiya, M., et al., 2016; Giuliani, A., et al, 2018]. Given CP acts as genome-attractor, it is essential to understand how the change in the CP leading to super-critical genome occurs. To elucidate the timing of the change in the CP, we focus on CM correlation dynamics (see **Methods**), i.e. the CM correlation between the initial and other time points: ***ĉ***^*k*^(*t*_0_). ***ĉ***^*k*^(*t*_*j*_) (*k* = 1,2, .., *K*) over experimental point, *t*_*j*_, where ***ĉ***^*k*^(*t*_*j*_) is unit vector (unit length) of the *k*^th^ group vector, ***c***^***k***^(*t*_*j*_). CM grouping is used to examine singular behavior at the CP.

#### 1. MCF-7 Breast Cancer Development (cell-population level): ON-OFF State of the CP Revealed

In MCF-7 breast cancer cells, the activation of ErbB receptor by HRG and EGF elicits two different biological responses. HRG-stimulation induces cell-differentiation, whereas EGF-stimulation provokes cell-proliferation [Nagashima T, et al., 2007; Nakakuki T, et al., 2010]. The temporal CM correlation in HRG stimulated MCF-7 cells reveals a divergent behavior at *t*_j_ = 15min (**Figure 4A**: left panel), whereas EGF-stimulated MCF-7 cells (right panel) do not show any divergent behavior. Both responses exhibit sandpile CPs (refer to Fig. 5A in [Tsuchiya, M, et al., 2016]). These results suggest that **the CP possesses both an activated and inactivated state**, i.e., ON/OFF expression states for a set of genes (**critical gene set**) corresponding to the CP. In EGF-stimulated MCF-7 cells, the CP is in the inactivated (OFF mode) state, whereas in HRG-stimulated MCF-7 cells, the CP is ON at 10-15min and thereafter turns OFF.

Direct evidence of ON/OFF state of the CP is shown as follows:

1. In **Figure 4B**, for HRG-stimulation, the switching of singular behaviors at 15-20min occurs at the CP. At the boundary of the CP (*ln*<*nrmsf*> ∼ -2.5), the singular behavior exhibits bimodal behavior in the fold-change on the CM grouping. At 15min, a dominant positive (i.e., fold-change greater than one) singular behavior (*ln*<*nrmsf*> > -2.5) suggests that swelled coil state of DNA occurs at the CP, while a negative singular behavior (*ln*<*nrmsf*> < -2.5) suggests compact coil state of DNA. At 20 min, this bimodal behavior switches to a dominant negative singular behavior (> -2.5) with a positive singular behavior (< -2.5). With regard to EGF-stimulation, such switching transition does not occur during the early time points. Note: a negative fold-change occurs due to the fact that the reference point is the CM of the whole expression. The subtraction of the CM(*t*_*j*_) from each gene expression has negative expression.
2. In **Figure 4C**, probability density function (PDF) of the whole expression [Shu G, et al., 2003] shows that at 15min, around the CP region, maximum of probability density occurs with positive value, whereas it becomes negative at 15-20min. This validates that the CP is in ON state at 15min and OFF state at 20min. PDF clearly shows that the ON-OFF switching of the CP induces global avalanche in whole gene expression.

It is worth noting that fold changes within expression groups, <***c***^*k*^(t_j+1_)>/<***c***^*k*^(t_j_)> exhibit a clear first-order phase transition involving genome sized DNA molecules (see more in **Discussion**). Through our studies, it became evident that the transition occurs as coherent behavior (at mega bp scale on chromosome) emerging from stochastic expression (coherent-stochastic behavior [Tsuchiya, M., et al., 2017]). This is to say that the averaging behavior (mean-field) of group expression corresponds to the coherent transition. On the other hand, ensemble average of fold change in individual expressions between two temporal groups, <***c***^*k*^(t_j+1_)/***c***^*k*^(t_j_)> does not reveal such characteristics in the transition. Interestingly, ensemble average of time difference in the expression group, <***c***^*k*^(t_j+1_)- ***c***^*k*^(t_j_)> supports the coherent scenario. This indicates that fluctuation (noise) on coherent dynamics is eliminated (see attached **Supplementary Figure S1**).

Regarding cell-fate change in HRG-stimulated MCF-7 cells, the erasure of the initial-sandpile criticality occurs after 2h (**Figure 4D**). While the cell-fate guiding critical transition occurs at an earlier time point (15-20min), the cell fate change happens after 2h. This time lag is needed to develop a new cell fate attractor through the coordinated local chromatin interaction (see details in [Zimatore, G., et al., 2019]).

#### 2. HL-60 Breast Cancer Cell Development (Cell-Population Level): Timing of the Cell-Fate Change

Cell development in HL-60 human leukemia cells further supports the scenario: 1) State of the CP changes such as from inactivate to activated state (OFF-ON) or ON-OFF while 2) the switching of singular behaviors occurs and 3) induces cell-fate guiding global avalanche. As for the switching behaviors at the CP,

A. In atRA-stimulation (**Figure 5B**), the dominant negative fold-change (compact globule state) indicates OFF-state of the CP at 18-24h, while the dominant positive fold-change (swelled coil state) for ON-state at 24-48h: OFF to ON state occurs at the CP. The global avalanche is shown by swelling out vivid probability density from 18-24h to 24-48h and contracting it at 48-72h (**Figure 5C**).
B. In DMSO-stimulation (**Figure 6B**), the dominant positive to negative fold-change (reverse to the atRA-stimulation) occurs from 8-12h to 12-18h, i.e., OFF to ON state occurs at the CP. At 18-24h, the CP is neither ON or OFF. These are confirmed in temporal change in the PDF profile (**Figure 6C**) around the CP: the fold-change (*ln*<*nrmsf*> ∼ -2.5) from the positive to negative to neutral.

**Figure 6:**
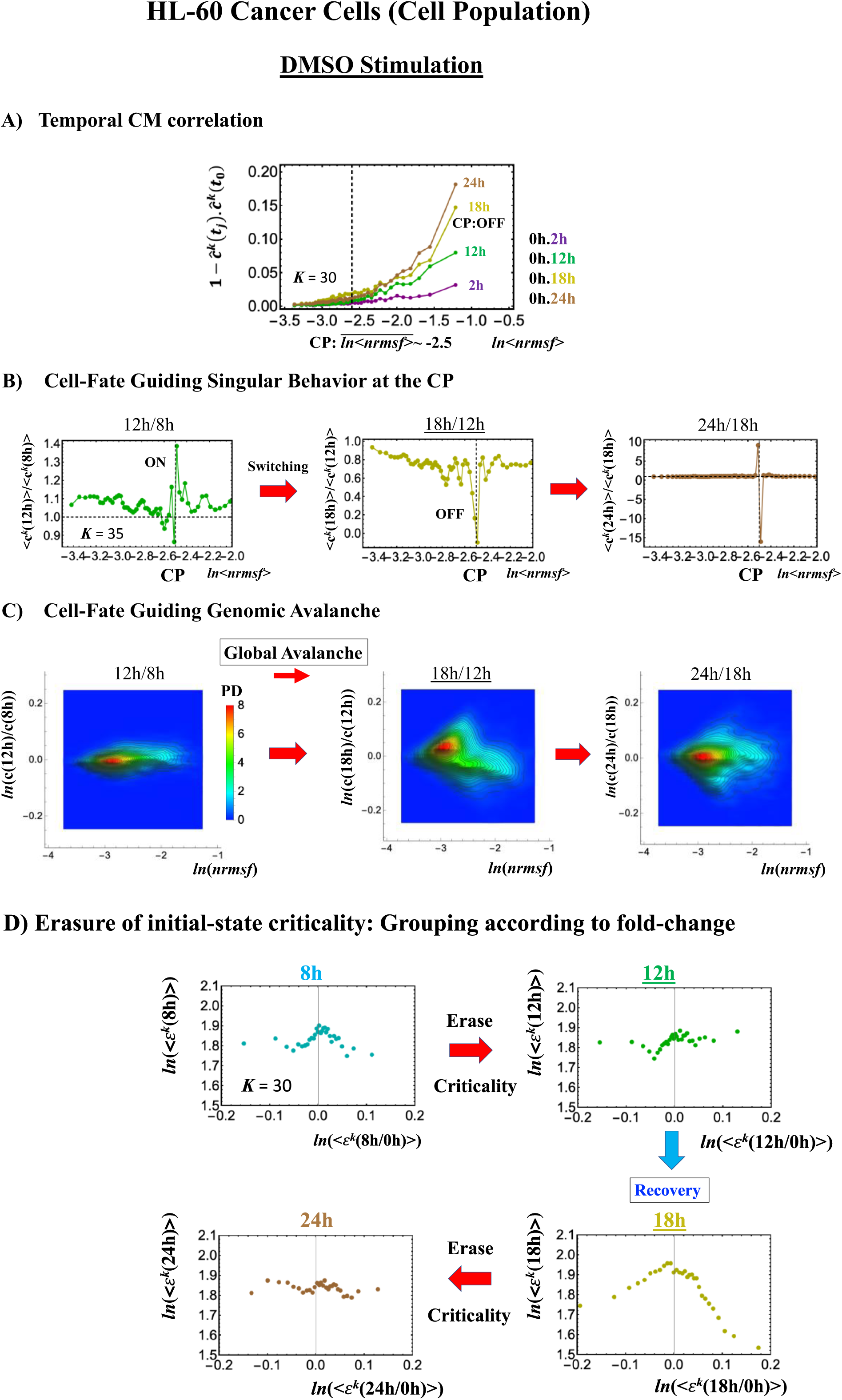
DMSO-stimulated (right) HL-60 cell population. **A)** Deactivation (OFF) of the CP occurs at12-18h for DMSO as shown by the temporal CM correlation response (*K* = 30: *n* = 315 mRNAs). The positive response occurs for OFF-state at 18h. This is due to the cosine function (even-function around zero) of two expression vectors (**Methods**). **B)** Transition of the higher-order structure at the CP (*K* = 35: *n* = 360 mRNAs) from swelled coil (ON) to compact globule state (OFF) to neutral state (neither ON or OFF; see **C**) at 18-24h is suggested to occur at 12-18h and 18-24h for DMSO stimulation, when the initial-state criticality is erased (**D**). The sensitivity according to the number of groups is observed. **C)** Temporal change in PDF shows that probability density around the CP (*ln*<*nrmsf*> ∼ -2.5) change from positive to negative to zero, supporting results of **A)** and **B)**. This shows that global avalanche occurs at 12-18h and 18-24h. **D)** Full erasure happens at 8-12h and 18-24h for DMSO. The erasure of the initial-state criticality occurs at 12-18h and thereafter, recovering the initial-state criticality at 18h indicates the existence of two SOC landscapes at 8-12h and 12-18h, respectively (see more in [Tsuchiya, M., et al, 2017]).

Regarding the cell-fate change, the timing of activation of the CP through the switching of singular behaviors at the CP coincides with the timing of the erasure of the initial-state sandpile: at 24-48h for atRA stimulation (**Figure 5**) and at 12-18h for DMSO stimulation (**Figure 6**). This suggests that the cell-fate change occurs at 24-48h for atRA and 12-18h for DMSO.

As for the DMSO response, it is interesting to observe a multi-step process of erasure of initial-state sandpile criticality (**Figure 6D**): erasure at 8-12h; recovery of the criticality at 12h-18h; then erasure again at 18-24h. Multiple erasures suggest that the cell-population response passes over two SOC landscapes [Tsuchiya, M et al., 2016] at 8-12h and 18-24h.

The results obtained on cancer cells suggest that **the activation of critical gene set (CP) as the genome-attractor plays an important role in cell-fate change.**

### D. Cell-Fate Change in Single-Cell Dynamics

#### 1. Human Embryo Development: Genome Avalanche Along Lowly Expressed Genes

Embryo development starts from a single cell (zygote) and thus corresponds to a completely different situation with respect the cases discussed above, where the data refer to averages over millions of cells in a plate (i.e., cell population level). The CP acts as genome attractor at the single-cell level and in cell populations (**Figure 3**). In the temporal CM correlation, the CP is a point with no differential (**Figures 7A, 8A, 9A**), while it appears as a divergent point in cell populations. This feature reveals distinct response domains (critical states) in the single-cell genome expression.

**Figure 7:**
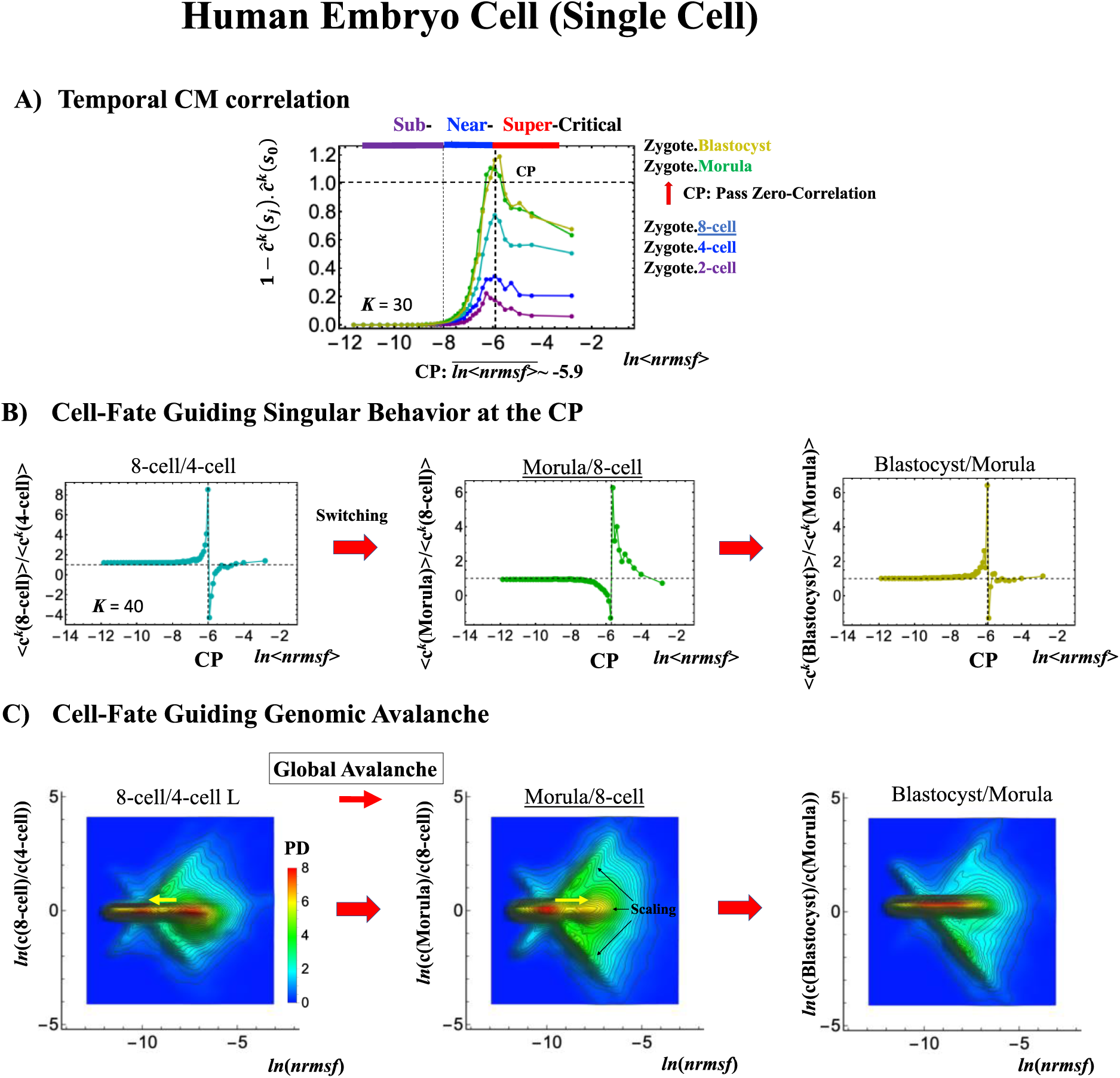
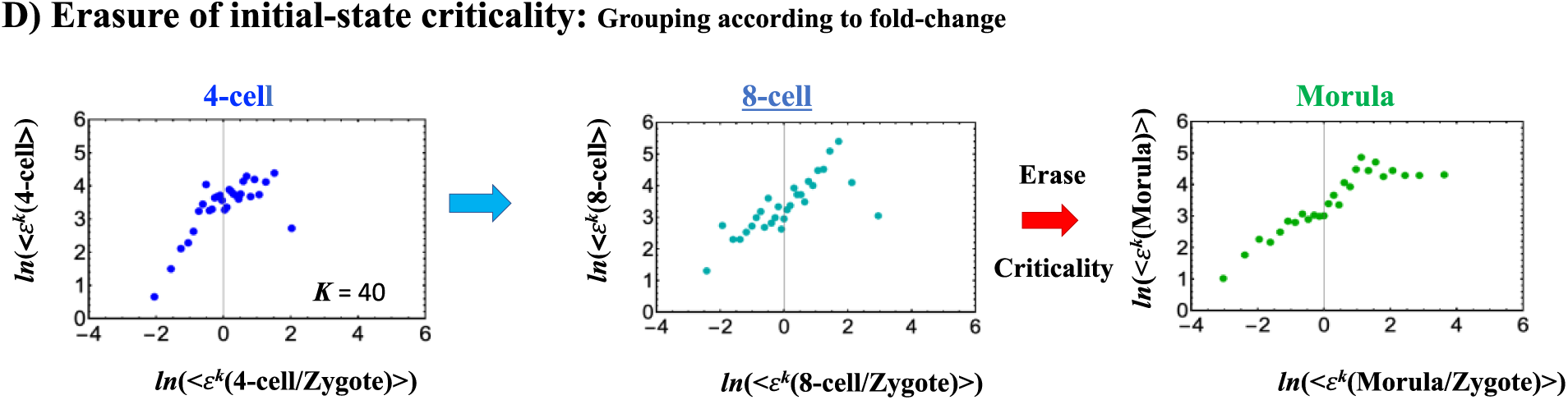
Human embryonic development (single cell). The CP corresponds to **A)** the peak of temporal CM correlation (see the difference from cell population in **Figures 4A, 5A**). A complete erasure of zygote-CP (genome-attractor) occurs after the 8-cell state (*K* = 30: *n* = 555 RNAs). Distinct response domains (critical states) as manifested in temporal CM correlations are shown (red: super-critical; blue: near-critical; purple: sub-critical), where location of the CP lies at the boundary between near- and super-critical states. **B)** Switching singular transition at the CP (*K* = 40: *n* = 416 RNAs) occurs at each cell state change (only shown from 4-cell here). The bimodal folding-unfolding feature at the CP suggests the occurrence of intra-chain segregation in the transition for genome sized DNA molecules (see more in **Discussion**). **C)** Genome avalanche occurs as the travelling density wave along lowly expressed genes (around non-fold change: *y* = 0; yellow arrow indicating the next travelling direction). Furthermore, three distinct scaling behaviors (linear density profiles in log-log plot) emerge, revealing that coordinated chromatin dynamics emerge to guide reprogramming. **D)** Erasure of zygote-criticality after the 8-cell state coincides with the timing of **A)** the zero temporal CM correlation through **B)** switching singular transition at the CP. This timing further matches with the timing of coherent perturbation on the genome-engine passing through the non-equilibrium fixed point (**Figure IIG**). Note: A linear behavior emerges in the grouping of randomly shuffled whole expression as expected; therefore, higher nrmsf is associated with higher expression (see more in Fig. 3 [Tsuchiya, M., et al., 2016]).

**Figure 8:**
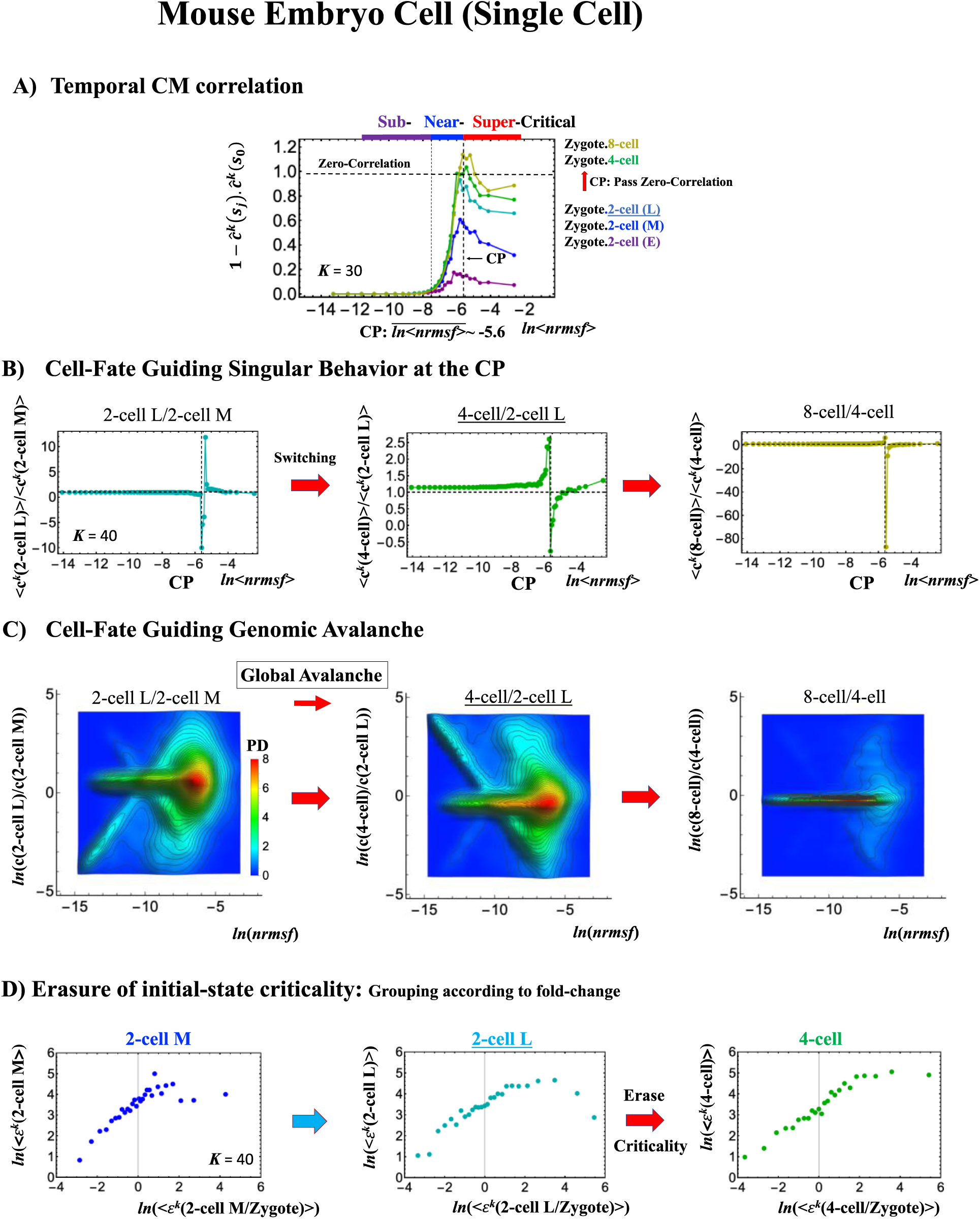
Mouse embryonic development (single cell). **A)** The complete erasure of zygote-CP (*K* = 30: *n* = 571 RNAs) occurs after the late 2-cell state **B)** through the switching singular transition at the CP (*K* = 40: *n* = 428 RNAs). **C)** Cell-fate guiding genome avalanche occurs from the middle 2-cell (2-cell M) to 4-cell state through the late 2-cell state (2-cell L), where the whole PDF profile reflects along the zero-fold change (*y*-axis) to exhibit coherent switching of folding and unfolding chromatin dynamics. The CP is ON at the middle 2-cell - late 2-cell state, OFF at the late 2-cell - 4-cell state and at the 4-cell state - 8-cell state. **D)** Erasure of initial-sandpile criticality occurs after late 2-cell state, coinciding with the result of **A)** and confirming the timing of cell fate change (after the late 2-cell state). Note: the travelling wave is also observed as shown in the human embryo cell (**Figure 7C**) between the zygote and other states (data not shown).

**Figure 9:**
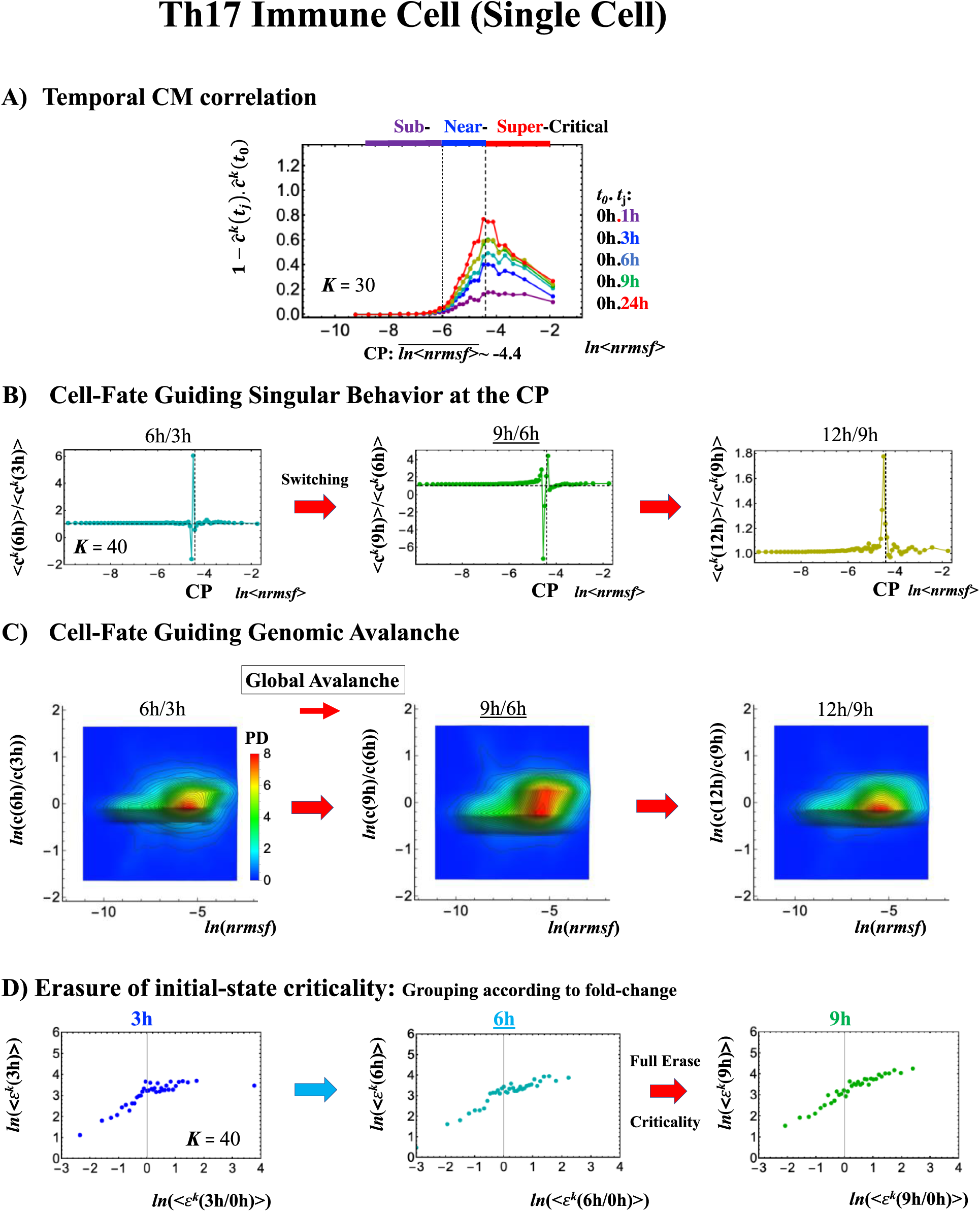
Th17 immune cell development (single cell). **A)** The CP features are the same as the embryo cases, with the only difference that the CP never passes the zero-temporal correlation (*K* = 30, n = 437 RNAs). **B)** The switching singular transition (*K* = 40: n = 328) at the CP (genome-attractor) occurs from 3h to 9h through 6h, which induces **C)** large-scale (not whole genome as in the embryo cases) expression avalanche: at 6-9h swelling out the density profile from 3-6h and then, contraction at 9-12h. The timing of cell-fate change after 6h is demonstrated by **D)** erasure of the initial-state (*t* = 0) criticality. This timing is further supported by the same timing occurrence as seen in the switch from suppression to enhancement on the genome engine (**Figures 13E**). The sensitivity of singular transition is observed according to the number of groups.

In the reprogramming event, the temporal CM correlation for the CP traverses zero value (corresponding to random-like behavior) at the 8-cell state for a human embryo cell (**Figure 7A**). This implies that after the 8-cell state, the human embryo cell completely erases the initial zygote criticality. Furthermore, this coincides with the erasure of the sandpile-type zygote CP (**Figure 7D**). In biological terms, this corresponds to the erasure of the initial stage of embryogenesis (driven by maternal heredity; see “SOC Control in Human and Mouse Related to the Developmental Oocyte-to-Embryo Transition” in Discussion [Tsuchiya, M., et al, 2016]).

It is important to note that groups of low-*nrmsf* presenting flattened CM correlations in time (*ln*<*nrmsf>* **<** -8.0) do not point to a no-response situation; on the contrary, they behave in a highly coherent manner to generate the autonomous SOC mechanism (see e.g., Fig.6 in [Tsuchiya, M, et al., 2015]).

The switching transition of the CP (**Figure 7B**) occurs at every cell state change from zygote to blastocyst (only results from 4-cell to blastocyst are shown). This suggests that in early embryogenesis, change in cell state involves a global change (genome avalanche) in whole expression. Notably, temporal change in PDF profile shows that genome avalanche occurs along lowly expressed genes (**Figure 7C**: around zero value in *y*-axis) shown as traveling density wave (higher to lower *nrmsf*) from the 8-cell to the morula state. The reverse travelling wave (lower to higher *nrmsf*) occurs after the morula state. This travelling wave points to the import role of collective behaviors emerged in lowly expressed genes (refer to local sub-critical state as the generator of genome-engine in **section IIF**). It is intriguing to observe that scaling behaviors (in log-log plot: **Figure 7C**) emerges, where most vivid scaling behaviors develop at the 8-cell - morula state.

Therefore, with the result of temporal CM correlation, we conclude that a major cell-fate change (embryonic reprogramming) occurs after the 8-cell state. This coincides with the timing of inverse coherent perturbation on the genome engine (cyclic flux flow), where the genome system passes over the non-equilibrium fixed point after 8-cell (onset of symmetry breaking occurs; see later section).

#### 2. Mouse Embryo Development: Switching Scaling Behavior in Cell-Fate Change

In mouse embryo development, the scenario of cell-fate change described above is confirmed:

1. Complete erasure of the memory of zygote CP occurs after the late 2-cell state (**Figure 8A**), where the switching singular transition occurs in cell-state change from the middle 2-cell (2-cell M) to the 8-cell state (**Figure 8B**), showing the major change in the genome.
2. This is confirmed by temporal change in PDF (**Figure 8C**): During cell development from the middle 2-cell to 4-cell state through the late 2-cell state (2-cell L), the whole density profile reflects along the axis of zero-fold change. This clearly manifests as the ON state of the CP at the middle 2-cell - late 2-cell state, OFF at the late 2-cell - 8-cell state. There are also three distinct scaling behaviors as shown in human embryo development. Notably, during the middle 2-cell to 4-cell through the late 2-cell state, the reflection of the scaling behaviors along the zero-fold change axis illustrates that in the genome-wide scale, coherent switching of folding and unfolding chromatin dynamics occurs.
3. The timing of cell-fate change is further confirmed by the erasure of zygote sandpile criticality after the late 2-cell state.

#### 3. Th17 Immune Cell Development: Partial Avalanche Guides Cell-fate Change

Cell-fate change in Th17 immune single cell development again follows the same general scenario. Compared with embryo development, there are several distinct differences worth noting:

1. The CP does not pass over zero temporal correlation with the initial state (*t* = 0h) although the correlation response increases in time, which indicates that cell differentiation induces a partial-scale (specific set) change in the whole expression (vs. whole-scale change in embryonic reprogramming).
2. The switching singular transition at the CP (genome-attractor) occurs from 3h to 12h (**Figure 9B**). This manifests as the OFF state of the CP at 6-9h and ON at 9-12h. **Figure 9C** illustrates that the change in expression for the switching still involves a large-scale, which is confirmed by the temporal change in PDF of the whole expression. However, there is no distinct scaling behavior as in the embryo development.

The timing of cell-fate change, as in other biological regulations, is determined by the erasure of the initial-state sandpile criticality (**Figure 9D**). This appears at 6h where an inversion of singular behaviors of the CP takes place, which is further confirmed by the timing of the inversion of perturbation on the genome-engine (**Figure 13E**). The OFF-ON switching at the CP guides the cell-fate change.

**Figure 10:**
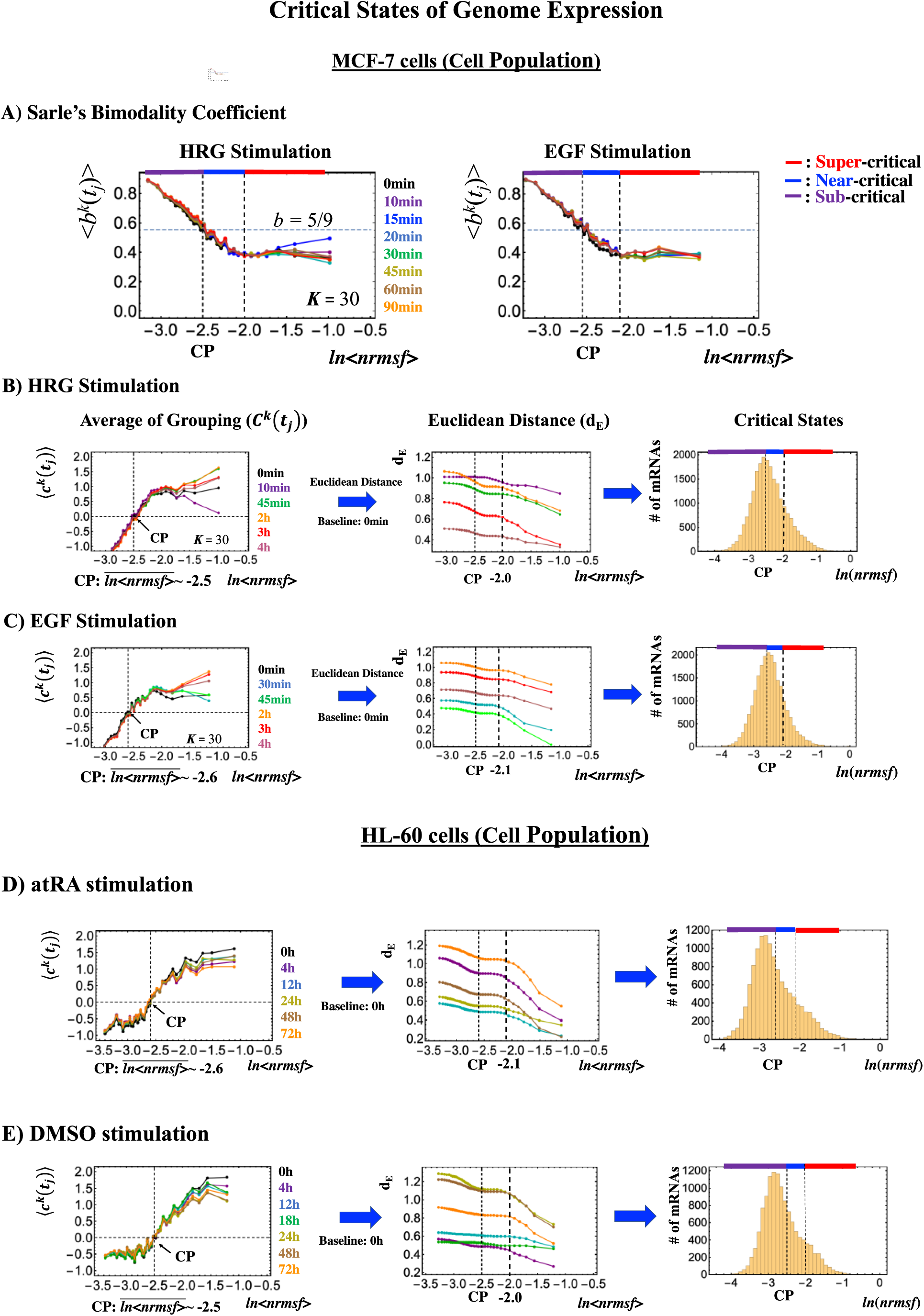
Systemic determination of critical states for cell-population (microarray data). **A)** In MCF-7 breast cancer cells, Sarle’s bimodality coefficient (*b* >5/9: onset of bimodal profile) shows that around the CP, a unimodal to bimodal transition (from high to low *nrmsf*) occurs for the group expression profile (*K* = 30: *n* = 742 mRNAs), which reveals three distinct profiles (red: super-; blue: near-; purple: sub-critical state) in the whole expression (left: HRG-stimulation; right: EGF-stimulation). To confirm these distinct behaviors (see panels **B)** and **C)**), which is based on the temporal response of the CM group (first column; refer to **Figure 2**), we examine the Euclidian distance (second column) between two ensemble averages, <***c***^*k*^(*t* = 0) > and < ***c***^*k*^(*t*_*j*_) > (*k* = 1, 2,..,*K =* 25) from higher *nrmsf* and confirm three distinct behaviors with a boundary indicated by dashed vertical lines. Similarly, for HL-60 human leukemia cells corresponding results are shown in **D)** atRA-stimulated and **E)** DMSO-stimulation. In third column (**B**-**E**), the corresponding region of critical states is shown in a histogram of gene expression according to *ln*(*nrmsf*). **Note**: The location of the CP lies at the boundary between near- and sub-critical states, different from the case with single cell (**Figures 7, 8, 9**).

**Figure 11:**
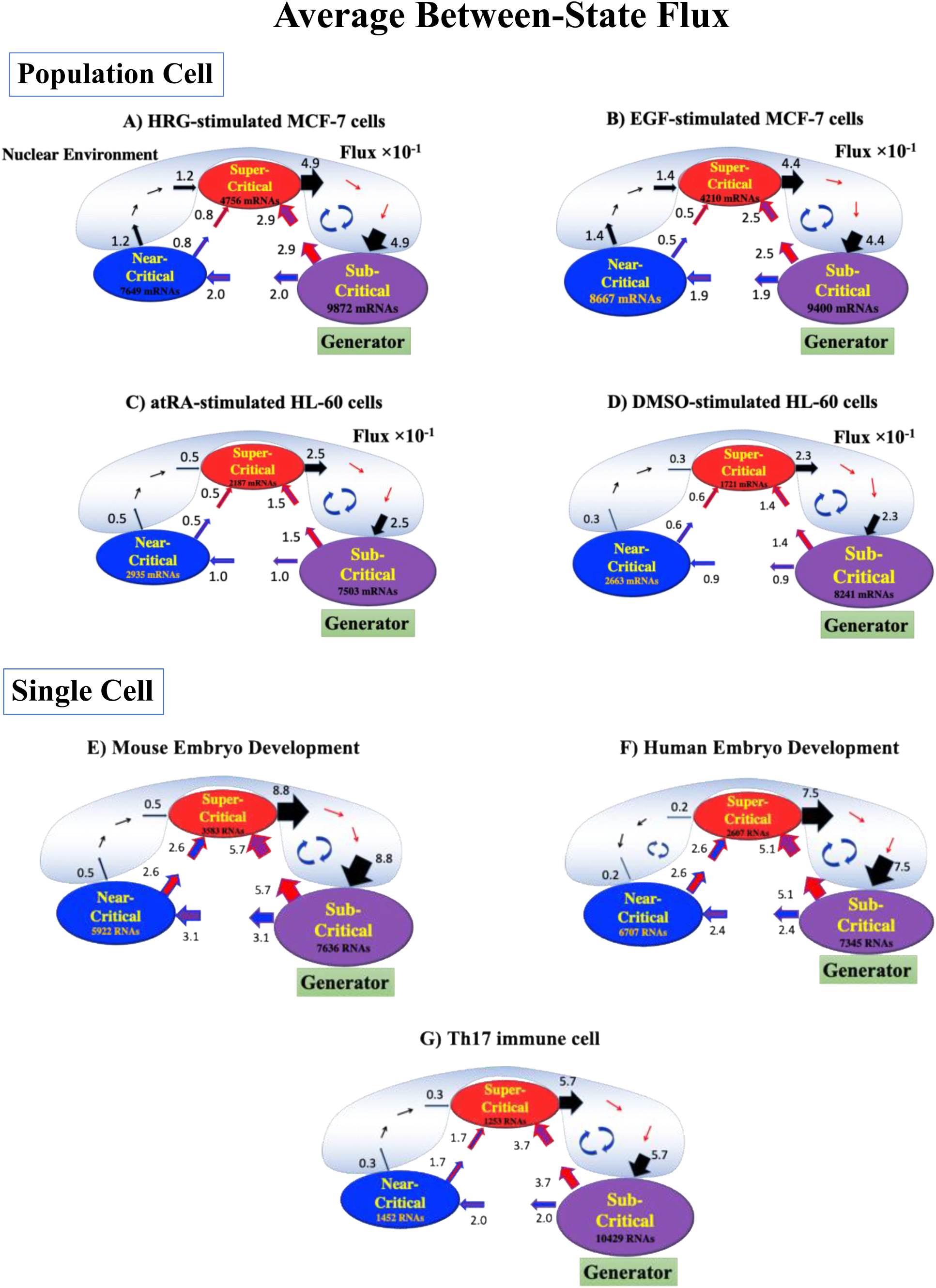
Genome engine mechanism revealed through average between-state expression flux. Cell-Population: **A)** HRG-stimulated and **B)** EGF-stimulated MCF-7 cancer cells; **C)** atRA-stimulated and **D)** DMSO-stimulated HL-60 cancer cells. Single-Cell**: E)** mouse- and **F)** human-embryo development; **G)** Th17 immune cell development. Figures show a common genome engine mechanism [Tsuchiya, M., et al., 2016]: Sub-super cyclic flux forms a dominant flux flow to establish the genome engine mechanism. The sub-critical state acting as a “large piston” for short moves (low-variance expression) and the super-critical state as a “small piston” for large moves (high-variance expression) with an “ignition switch” (a critical point: the genome attractor) are connected through a dominant cyclic state flux as a “camshaft”, resulting in the anti-phase dynamics of two pistons. Numerical values represent average between-state expression flux, whereas in the cell population, the values are based on 10^−1^scale.

**Figure 12:**
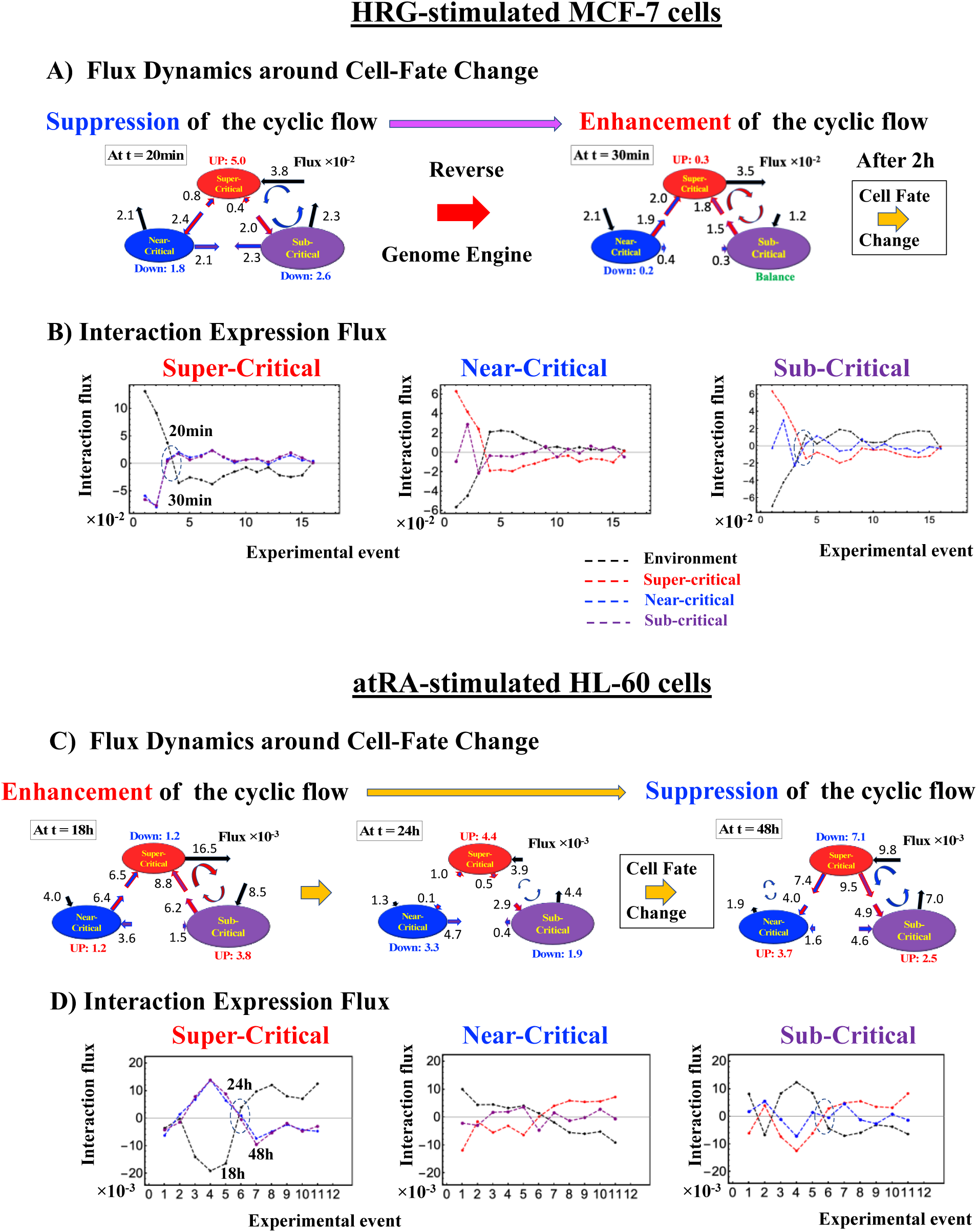
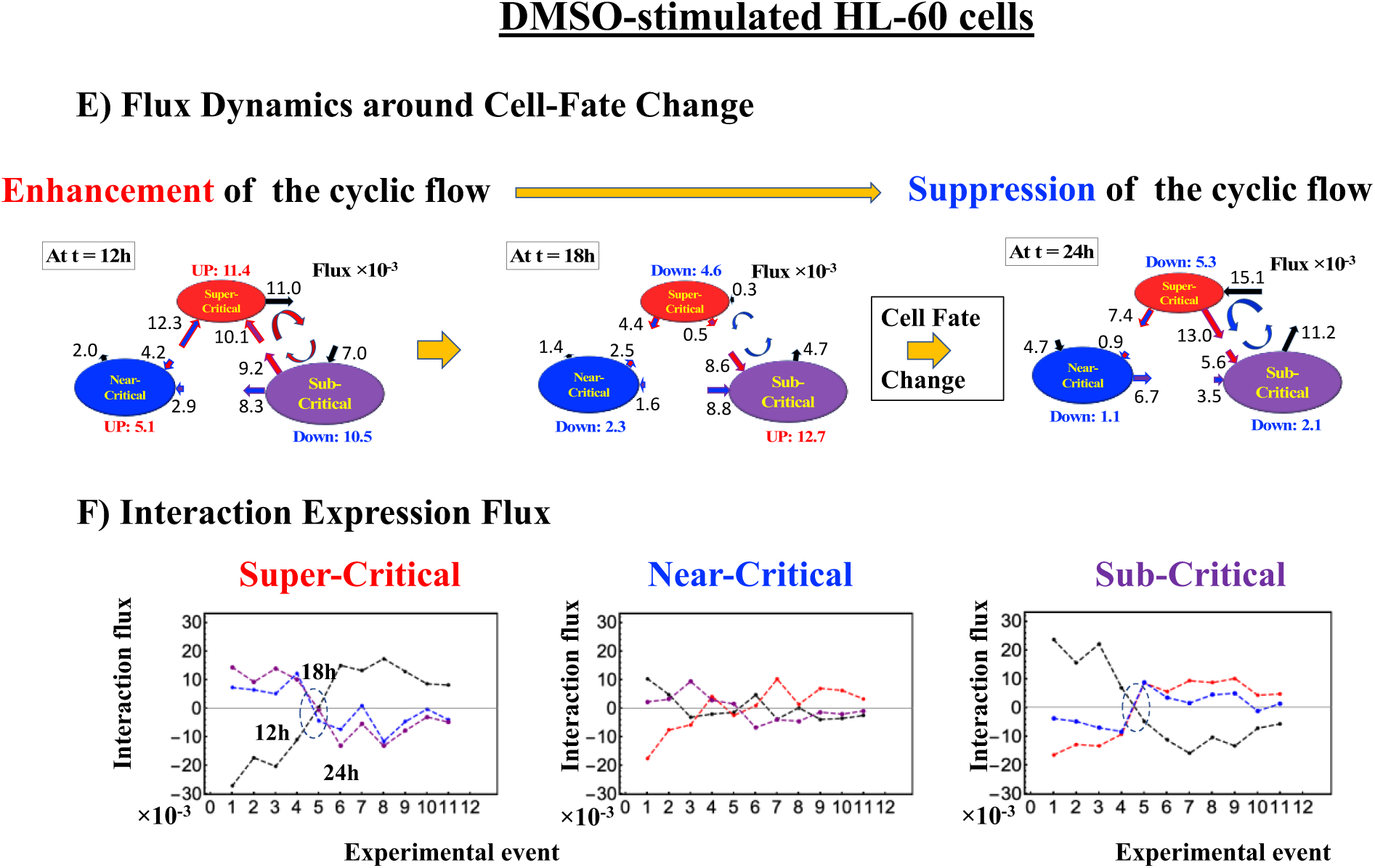
Cell-population - Cell fate mechanism through perturbation on the genome-engine. **A)** and **B)** MCF-7 cells**; C)** and **D**) HL-60 cells. Before the cell-fate change, the interaction flux dynamics (indicated by dashed cycle: **B, D, F**) intersect each other. This marks a non-equilibrium fixed point and the onset of the switch to coherent dynamics on the dominant cyclic flow (genome-engine: **Figure 11**). The interaction flux dynamics (black: environment; red: super-critical state; blue: near-critical state; purple: sub-critical state) show that on the dominant cycle state flux, **A)** suppression to enhancement for MCF-7 cells occurs before the cell-fate change (**Figure 4D**), **B)** and **C)** switching from enhancement to suppression for HL-60 cells during the cell-fate change. Numerical values based on 10^−2^ for MCF-7 cells and 10^−3^ scale for HL-60 cells represent interaction flux. Refer to **Methods** for further detail experimental time points.

**Figure 13:**
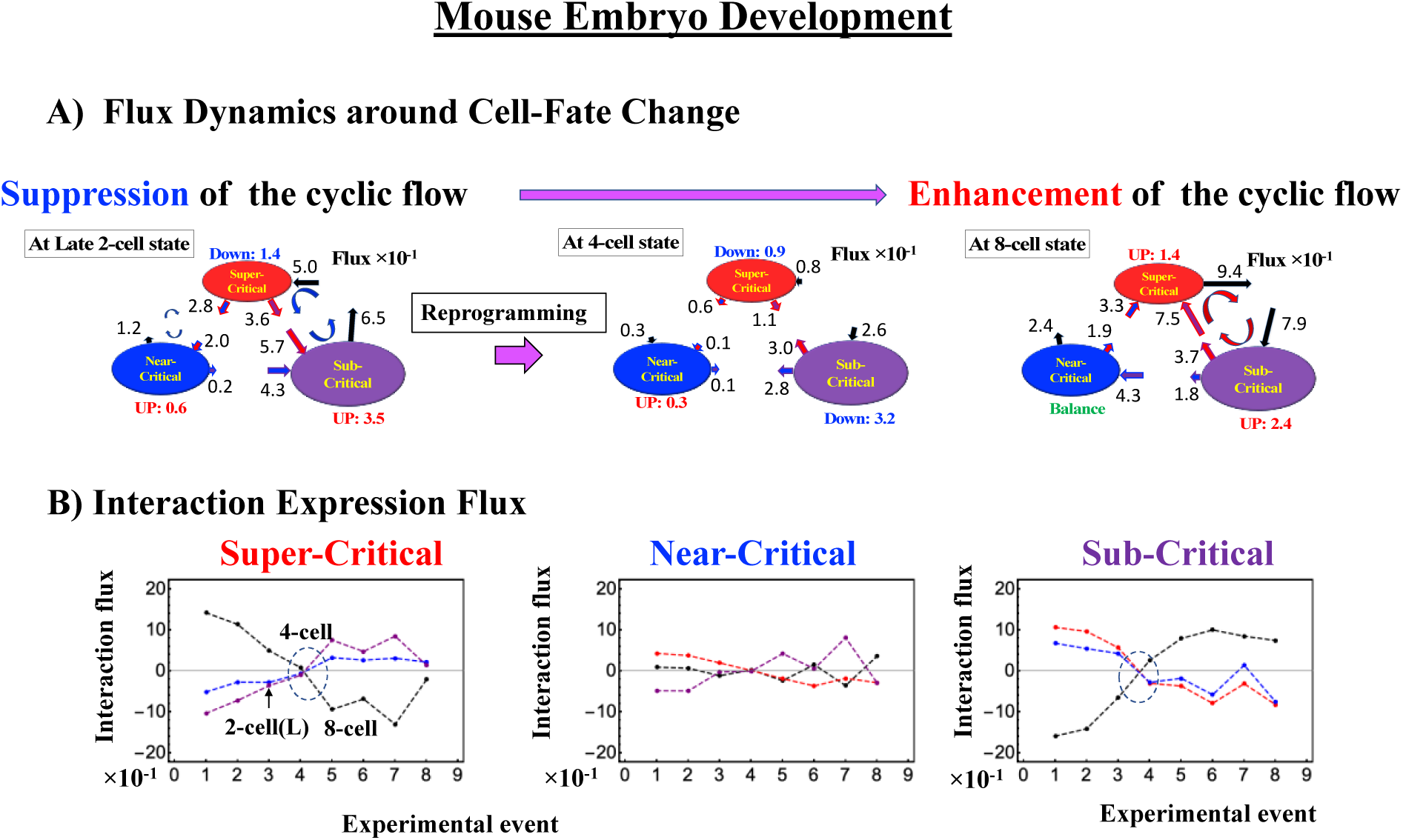
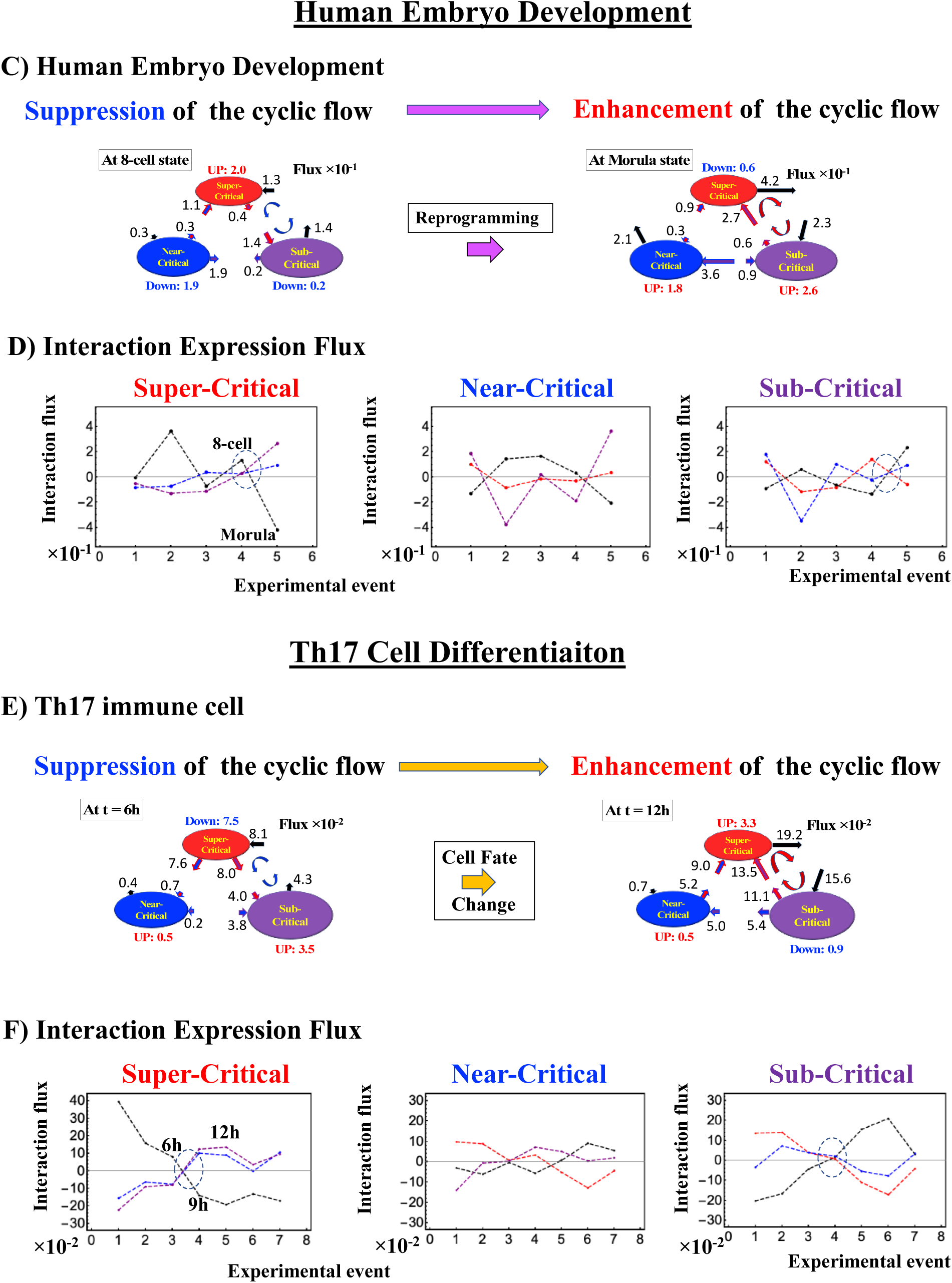
Single cell - Cell fate mechanism through perturbation on the genome engine. **A)** and **B)** Mouse embryo development**; C)** and **D)** Human embryo development; **E)** and **F)** Th17 immune cell differentiation. Notably, reprogramming on embryo development (more evident on mouse embryo) and cell differentiation on Th17 immune cell occurs through the suppression-enhancement on the dominant cycle state flux between super- and sub-critical states, which is opposite to the cell differentiation in HL-60 cells; for HRG-MCF-7 cell response refer to Fig. 12B in [Tsuchiya, M., et al., 2016]. Numerical values based on 10^−1^ for embryo cell and 10^−2^ scale for Th17 cell represent between-state expression flux.

Cancer cell-differentiation goes together with compact globule (OFF) to swelled coil (ON) or vice versa at the CP (see **Figures 4-6**). As shown in **Figures 5C-6C** (HL-60 cells), the global genome avalanche occurs at the timing of cell-fate change. As for MCF-7 cells (HRG-stimulation), the timing of cell-fate change (after 2h: **Figure 4D**) occurs later than that of the genome avalanche (see detail attractor mechanism in [Zimatore, G. et al, 2019]). The global impact provoked by the CP is further supported by the fact that 1) EGF-stimulated MCF-7 cells, where the CP is OFF with no cell-differentiation, induce only local perturbation (**Figure 14C**) and 2) HL-60 cells coincides with the timing of global perturbation (**Figure 14D**).

Regarding cell-fate change on single cell dynamics, a global perturbation (**Figures 14A, B**) occurs on the genome-engine (after the late 2-cell, 8-cell and 6h for mouse, human embryo, and Th17 cell, respectively), where the timing coincides with that of the cell-fate change (i.e., erasure of the initial-state CP memory). This suggests that the cell-fate guiding change in the CP corresponds to the erasure of initial-state sandpile criticality. As for embryo reprogramming, this picture is supported by the fact that temporal CM correlation of the CP from initial state (zygote) passes zero-correlation (i.e., erasure of the initial-state CP memory: **Figures 7A,8A**). Due to genome attractor, the change in the CP (the edge of the criticality) provokes a global impact on the entire genome expression through the genome-engine perturbation: the global perturbation leads to the occurrence of genome avalanche at the timing of cell-fate change (**Figures 7C-9C**; also refer to Fig.13 and Discussion in [Tsuchiya, M., et al., 2016]).

### E. Systematic Determination of Local Critical States in Genome Expression

We demonstrate that the CP is a fixed-point relative to a given biological regulation. Next, based on this fact, we show that critical states in genome expression can be determined systematically for both single-cell and cell-population genome expression:

1. **Single-cell level**: both temporal and spatial CM correlations (**Figures 7A,8A,9A**) manifest distinct response domains according to *nrmsf*: low-variance expression (sub-critical state) for the region of flattened correlation, intermediate-variance for the near-critical state from the edge of flattened correlation to the CP, and high-variance expression for super-critical state above the CP (summary: **Table 1**).

**Table 1:** Critical States of Single-Cell Genome Expression.
2. **Cell-population level**: As shown in **Figures 3,4**, the Euclidian distance (from the highest *nrmsf* group) between the temporal responses of CM grouping (**Figure 2**: *t* = 0 vs. other experimental time points) reveals critical states (**Figure 10**; summary: **Table 2**), where the CP exists at the boundary between near-and sub-critical states (vs. between super-and sub-critical states in single cell). This occurs for both MCF-7 and HL-60 cancer population cells. In our previous studies on microarray data (cell-population level), critical states were determined by the transition of expression profile by means of Sarle’s bimodality coefficient (**Figure 10A**). This was accomplished by putting in evidence of distinct response domains (super-, near- and sub-critical domains according to temporal variance of expression), which were evident in genome expression (see more in [Tsuchiya, M., et al., 2015]).

| Mouse Embryo Development | Human Embryo Development | Th17 cell differentiation |
| --- | --- | --- |
| **Critical States:** | **Critical States:** | **Critical States:** |
| **Super-critical:** 3583 RNAs<br>$-5.6^* < \ln\langle nrmsf \rangle$ | **Super-critical:** 2607 RNAs<br>$-5.9^* < \ln\langle nrmsf \rangle$ | **Super-critical:** 1253 RNAs<br>$-4.4^* < \ln\langle nrmsf \rangle$ |
| **Near-critical:** 5922 RNAs<br>$-7.5 < \ln\langle nrmsf \rangle < -5.6$ | **Near-critical:** 6707 RNAs<br>$-8.0 < \ln\langle nrmsf \rangle < -5.9$ | **Near-critical:** 1452 RNAs<br>$-6.0 < \ln\langle nrmsf \rangle < -4.4$ |
| **Sub-critical:** 7636 RNAs<br>$\ln\langle nrmsf \rangle < -7.5$ | **Sub-critical:** 7345 RNAs<br>$\ln\langle nrmsf \rangle < -8.0$ | **Sub-critical:** 10429 RNAs<br>$\ln\langle nrmsf \rangle < -6.3$ |

\*Critical Point at the boundary between **Super-** and **Near-**critical states

**Table 2:** Critical States of Cell Population Genome Expression.

| HRG-MCF-7 cells | EGF-MCF-7 cells | atRA-HL-60 cells | DMSO-HL-60 cells |
| --- | --- | --- | --- |
| **Critical States:** | **Critical States:** | **Critical States:** | **Critical States:** |
| **Super-critical:** 4756 mRNAs<br>$-2.0 < \ln\langle nrmsf \rangle$ | **Super-critical:** 4210 mRNAs<br>$-2.1 < \ln\langle nrmsf \rangle$ | **Super-critical:** 2187 mRNAs<br>$-2.1 < \ln\langle nrmsf \rangle$ | **Super-critical:** 1721 mRNAs<br>$-2.0 < \ln\langle nrmsf \rangle$ |
| **Near-critical:** 7649 mRNAs<br>$-2.5^* < \ln\langle nrmsf \rangle < -2.0$ | **Near-critical:** 8667 mRNAs<br>$-2.6^* < \ln\langle nrmsf \rangle < -2.1$ | **Near-critical:** 2935 mRNAs<br>$-2.6^* < \ln\langle nrmsf \rangle < -2.1$ | **Near-critical:** 2663 mRNAs<br>$-2.5^* < \ln\langle nrmsf \rangle < -2.0$ |
| **Sub-critical:** 9872 mRNAs<br>$\ln\langle nrmsf \rangle < -2.5$ | **Sub-critical:** 9400 mRNAs<br>$\ln\langle nrmsf \rangle < -2.6$ | **Sub-critical:** 7503 mRNAs<br>$\ln\langle nrmsf \rangle < -2.6$ | **Sub-critical:** 8241 mRNAs<br>$\ln\langle nrmsf \rangle < -2.5$ |

\*Critical Point at the boundary between **Near-** and **Sub-**critical states

For both single-cell and cell-population levels, the existence of distinct critical states is supported by the fact that 1) mixed critical state does not converge to the CM of the critical state due to distinguished coherent dynamics of critical states and 2) distinct stochastic behaviors are shown among critical states (see e.g., Figs. 3D and 3B in [Tsuchiya, M., et al., 2017] for single cell, respectively).

### F. Genome-Engine Mechanism for SOC-Control of Genome Expression

The co-existence of distinct local critical states with a fixed critical point shows that the whole genome expression is dynamically self-organized through the emergence of a critical point (CP) from embryo to cancer development. This points to the existence of an autonomous critical-control genomic system.

Distinct coherent dynamics in critical states emerge from stochastic expression and the CM of critical state represents its dynamics (Fig. 10 in [Tsuchiya, M., et al., 2016]). Thus, dynamic expression flux analysis [Tsuchiya, M., et al., 2016, 2018; **Methods**] to the CM of critical states can apply to reveal the genome-engine mechanism for describing how autonomous SOC control of genome expression occurs. **Figure 11** shows that the sub-critical state acts as internal source of expression flux and the super-critical state acts as a sink. The cyclic flux forms a dominant flux flow that generates a strong coupling between the super- and sub-critical states accompanied by their anti-phase expression dynamics. This results in making its change in oscillatory feedback to sustain autonomous SOC control of overall gene expression. Thus, average between-state flux (**Figure 11**) represents a stable manifold for the thermodynamically open system. The formation of the dominant cyclic flux provides a *genome-engine metaphor* of SOC control mechanisms pointing to a universal mechanism in gene-expression regulation of mammalian cells for both population and single-cell levels. Global perturbation stemmed from the change in the CP, which enhances or suppresses the genome-engine, ends up into cell-fate change.

### G. A Universal Genome-Engine Mechanism for Cell-Fate Change: CP as the Organizing Center of Cell-Fate

The key point of genome engine mechanism is that dynamics of distinct coherent behavior emerges from stochastic expression in local critical states (coherent-stochastic behavior) and follows the dynamics of the CM. The CM acts as attractor at both whole genome (**Figure 2**) and critical state levels (e.g., Fig. 3B in [Tsuchiya, M., et al., 2017]). Based on this fact, the expression flux (effective force) approach (**Methods**) was developed to reveal dynamic interaction flux between critical-state attractors (Fig. 10 in [Tsuchiya, M., et al., 2016] and Fig. 3 in [Tsuchiya, M., et al., 2017]). This is instrumental to demonstrate that a heteroclinic critical transition guides coherent behaviors emerged in the critical states. Interaction flux between-state attractors is the underlying basic mechanism of epigenetic self-regulation incorporating a rich variety of transcriptional factors and non-coding RNA regulation to determine coherent oscillatory behaviors of critical states.

The flux dynamics approach is further developed to analyze quantitative evaluation of the degree of non-harmonicity and time reversal symmetry breaking in nonlinear coupled oscillator systems [Tsuchiya, M., et al., 2018].

We can summarize the systemic view about how and when cell-fate change occurs, as follows:

1. **How cell-fate change occurs**: Intersection of interaction fluxes (**Figure 12** for cell population and **Figure 13** for single cell) occurs just before cell-fate change. This intersection means that thermodynamically, the genomic system lies in a non-equilibrium fixed point (e.g., see in **Figure 12C**: atRA-stimulation at 24h with small interaction fluxes; see more **Methods**), which suggests that before cell-fate change, time-reversal symmetry is broken through passing over a non-equilibrium fixed condition. In terms of genome-engine, around the cell-fate change, a global perturbation induces enhancement or suppression on the genome-engine, where there is a dominant cyclic flux flow between the super- and sub-critical states (**Table 3**: single cell; **Table 4**: cell population). In HL-60 cells (cell population), the genome-engine is enhanced before the cell-fate change and suppressed (enhancement-suppression) thereafter. On the contrary, a reverse process of suppression-enhancement takes place in the MCF-7 cancer cells (**Figure 12A**; see more in [Tsuchiya, M., et. al., 2016]). In the single-cell cases (embryo development and Th17 immune cell), a suppression-enhancement on the genome engine occurs (**Figure 13**). The varying sequences of perturbation on the genome-engine may stem from different stages of the suppressive pressure on cell-differentiation against cell-proliferation.

**Table 3:** Single Cell: State Change in the CP associated with Coherent Perturbation on the Genome Engine.

|  | Location of the CP | State-Change in the CP | Coherent Perturbation on Genome-Engine | Reprogramming*/ Cell fate change |
| --- | --- | --- | --- | --- |
| Mouse Embryo Development | $\ln\langle nrmsf \rangle_{CP} \sim -5.6$ | **ON** → **OFF**:<br>At late 2-cell - 4-cell<sup>#</sup> | Suppression-Enhancement | After late 2-cell* |
| Human Embryo Development | $\ln\langle nrmsf \rangle_{CP} \sim -5.9$ | **Switching**:<br>At 8-cell – morula<sup>#</sup> | Suppression-Enhancement | After 8-cell* |
| Th17 Immune Cell Differentiation | $\ln\langle nrmsf \rangle_{CP} \sim -4.4$ | **OFF** → **ON**:<br>After 6-9h | Suppression-Enhancement | After 6h |

<sup>#</sup>Cell-fate guiding state change in the CP.

**Table 4:** Cell Population: State Change in the CP associated with Coherent Perturbation on the Genome Engine.
2. **When cell-fate change occurs:** Cancer cell-differentiation goes together with compact globule (OFF) to swelled coil (ON) or vice versa at the CP (see **Figures 4-6**). As shown in **Figures 5C-6C** (HL-60 cells), the global genome avalanche occurs at the timing of cell-fate change. As for MCF-7 cells (HRG-stimulation), the timing of cell-fate change (after 2h: **Figure 4D**) occurs later than that of the genome avalanche (see detail attractor mechanism in [Zimatore, G. et al, 2019]). The global impact provoked by the CP is further supported by the fact that 1) EGF-stimulated MCF-7 cells, where the CP is OFF with no cell-differentiation, induce only local perturbation (**Figure 14C**) and 2) HL-60 cells coincides with the timing of global perturbation (**Figure 14D**).

|  | Location of the CP | State-Change in the CP | Coherent Perturbation on Genome-Engine | Cell Fate Change |
| --- | --- | --- | --- | --- |
| EFG-stimulated MCF-7 cells | $\ln\langle nrmsf \rangle_{CP} \sim -2.6$ | OFF state | Local Perturbation | NO |
| HRG-stimulated MCF-7 cells | $\ln\langle nrmsf \rangle_{CP} \sim -2.5$ | ON → OFF:<br>At 15-20min | Suppression-Enhancement | After 2h* |
| DMSO-stimulated HL-60 cells | $\ln\langle nrmsf \rangle_{CP} \sim -2.5$ | ON → OFF:<br>At 12-18h | Enhancement-Suppression | After 18h |
| atRA-stimulated HL-60 cells | $\ln\langle nrmsf \rangle_{CP} \sim -2.6$ | OFF → ON:<br>At 24-48h | Enhancement-Suppression | After 24h |

\*Refer to time-lag mechanism in [Zimatore, G. et al, 2019].

Regarding cell-fate change on single cell dynamics, a global perturbation (**Figures 14A, B**) occurs on the genome-engine (after the late 2-cell, 8-cell and 6h for mouse, human embryo, and Th17 cell, respectively), where the timing coincides with that of the cell-fate change (i.e., erasure of the initial-state CP memory). This suggests that the cell-fate guiding change in the CP corresponds to the erasure of initial-state sandpile criticality. As for embryo reprogramming, this picture is supported by the fact that temporal CM correlation of the CP from initial state (zygote) passes zero-correlation (i.e., erasure of the initial-state CP memory: **Figures 7A,8A**). Due to genome attractor, the change in the CP (the edge of the criticality) provokes a global impact on the entire genome expression through the genome-engine perturbation: the global perturbation leads to the occurrence of genome avalanche at the timing of cell-fate change (**Figures 7C-9C**; also refer to Fig.13 and Discussion in [Tsuchiya, M., et al., 2016]).

Therefore, **the elucidation of activation-deactivation mechanism for the CP (involving more than hundreds of genes) through a coordinate chromatin structure change [**Zimatore, G. et al, 2019**] uncovers how the genome-engine is either enhanced or suppressed around the cell-fate change. Furthermore, our SOC hypothesis is expected to predict when and how a cell-fate change occurs in different situations (e.g., cancer, iPS cells, stem cells, etc.). The universality of the proposed model does not stem from our experimental results, but from the existence of very basic physical constraints (e.g., chromatin dimension and general organization, phase transition phenomenology) independent of biological specificities.**

**Figure 14:**
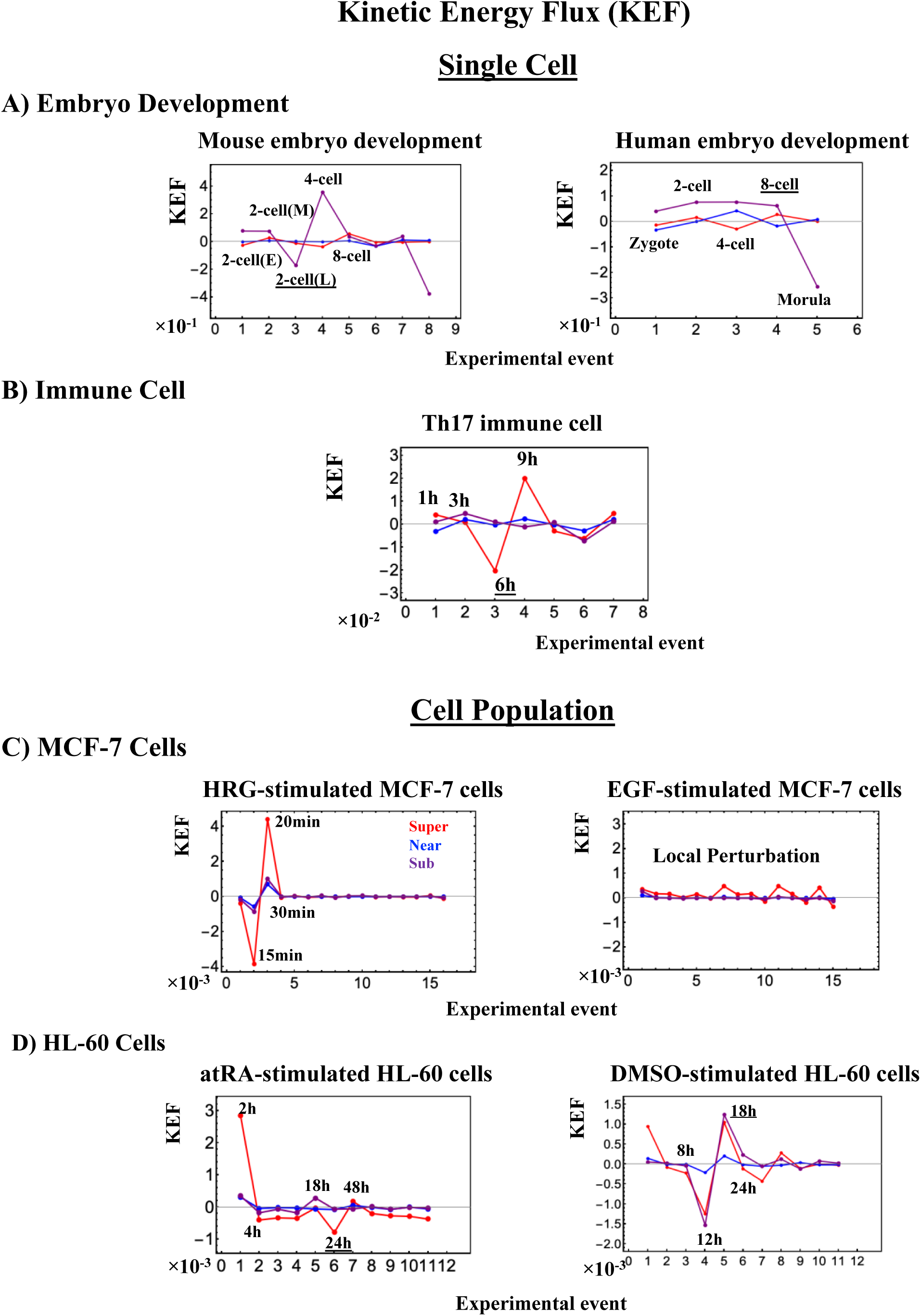
Kinetic Energy Flux (KEF). Global perturbation on genome-engine is well-manifested in the kinetic energy flux (**Methods**) examining single-cell responses in **A)** Embryo development and **B)** Immune cell, and population responses in **C)** MCF-7 cells and **D)** HL-60 cells. At the single-cell level, the timing of global perturbation, the involvement of more than one critical state, coincides with that of cell-fate change (**Figures 7C, 8C, 9C**).

At the population level, kinetic energy flux is damping. In the HRG-stimulated MCF-7 cells (**C**), the timing of the activation of the CP (ON at 15min; **Figure 4A**: left panel) is within the global perturbation (10-30min) (intersection of interaction flux at 20-30min in Fig [Tsuchiya, M, et al., 2016], whereas in the EGF-stimulation (right panel; no cell-differentiation [Saeki, Y., et al., 2009]), only local perturbation occurs. On the other hand, in HL-60 cells (**D**), the timing of the perturbation on the genome engine (at 18h-24h-48h intervals) corresponds to the end of the damping (second largest perturbation) for atRA-stimulated HL-60 cells (left panel) and the largest global perturbation (at 12h-18h-24h intervals) for DMSO-stimulated HL-60 cells (right panel). Underlined numbers represent the timing just before the cell-fate change.

## III. Discussion

In this study, we demonstrated the existence of a common underlying genomic mechanism for cell-fate decision holding from embryo to cancer development (**see Figure 1**). We summarize our findings in the following points:

1. **The critical point (CP) acts as the center of cell-fate**: The CP (a specific critical gene set) is (nearly) a fixed point according to temporal expression variance (*nrmsf*). The CP acts as the genome-attractor and its state change induces the global avalanche effect on genome expression (super-critical genome; see below) and determines the cell-fate change.
2. **How cell-fate change occurs**: Before cell-fate change, the genomic system passes over the stable point of the thermodynamically open system. The genome-engine, an emergent dynamic property of autonomous interaction flux between local critical states (distinct expression domains in whole expression) is enhanced or suppressed (**Tables 3, 4**) to induce coherent perturbation on the dominant cyclic flux between local super- and sub-critical states.
3. **When cell-fate change occurs:** Cell-fate change occurs at the timing of erasure of initial-state sandpile CP. At this time, the global genome avalanche occurs except for the HRG-stimulated MCF-7 cell differentiation, where there is an existing time-lag between the genome avalanche and cell-fate change (see further details for this mechanism in [Zimatore, G. et al, 2019]).

The evidence presented above provides a systematic determination of critical states with CP for both cell-population and single-cell regulation (**Tables 1, 2**). This implies that expression flux analysis among critical states through the cell nuclear environment provides a potential universal model of self-organization, leading to the emergence of a ‘genome-engine’. This ‘engine’ can be intended as a highly coherent behavior of low-variance genes (sub-critical state) entering into a dominant cyclic expression flux with high-variance genes (super-critical state) to develop autonomous critical control system. This explains the coexistence of critical states through the CP, with *nrmsf* acting as the order parameter for the self-organization.

CP shows a clear bimodal expression distribution reminiscent of swelled coil (ON) and compact globule (OFF) DNA transition corresponding to the inversion of the intra-chain phase segregation of coil and globule states. This behavior closely resembles the intrinsic characteristics of the first-order phase transition inherent in genome sized DNA molecules. The activation-deactivation of the CP is essential for the occurrence of cell-fate change.

As for mega base-pairs size of DNA phase transition, up until the late 20th century, it had been believed that a single polymer chain, including DNA molecule, always exhibit cooperative but mild transition between elongated coil and compact globule, which is neither a first-order nor a second-order phase transition [Flory, P., 1953; Gennes, P.G. 1979]. More recently, it has become evident that long DNA molecules above the size of several tens of kilo base-pairs exhibit characteristics to undergo large discrete transition, i.e., first-order property on coil-globule transition [Yoshikawa, K, et al., 1996; Yoshikawa, K and Yoshikawa, Y., 2002; Zinchenko A, et al., 2008]. Such first-order characteristics are rather general for a semi-flexible polymer, especially polyelectrolyte chains such as giant DNA molecules. Additionally, it is important to note that insufficient charge neutralization on the globule state causes instability and leads to the generation of intra-chain segregation on an individual single genomic DNA [Sakaue, T., Yoshikawa, K. 2006; Shew, C.Y., Yoshikawa, K., 2007] (see e.g., **Figure 8B**). When such instability with long-range Columbic interaction is enhanced, the characteristic correlation length tends to become shorter, corresponding to the generation of the critical state in the transition. Such behavior accompanied with the folding transition of DNA is also observed for reconstructed chromatin [Schiessel, H., 2003; Nakai, T., et al., 2005; Suzuki, Y., et al., 2011].

In relation to the classical concept of SOC (**Introduction**), the activation and inactivation of the CP suggests that there may be another layer of a macro-state (genome state) composed of distinct micro-critical states. When the state of the CP changes (i.e., change in the genome-attractor occurs), the perturbation can spread over the entire system in a highly cooperative manner, i.e., the genome-state to be considered ‘super-critical’ to determine the cell-fate change. Whereas, the state of the CP does not change in EGF-stimulated MCF-7 cells, where the genome states are considered *‘*sub-critical’.

Regarding genome-reprogramming, our results provide further insight of the reprogramming event in the mouse and human embryo development [Tsuchiya, M., et al., 2017, 2018]. In the mouse embryo development, energy release at the CP from ON to OFF state guides the genome-engine from a suppressed to enhanced state, and this drives the genome to pass over a critical transition state (SOC landscape) right after the late 2-cell state (note: regarding critical state transition in single-cell level, also see [Mojtahedi, M., et al., 2016]). The genome-engine suggests that the activation-deactivation mechanism of the CP should elucidate how the global perturbation occurs on self-organization through change in signaling by external or internal stimuli into a cell. Notably, around genome-reprogramming (human: the 8-cell state and mouse: the late 2-cell state), scaling behaviors in expression dynamics become apparent. This tells that coordinated chromatin dynamics emerge to guide reprogramming through SOC control. Recent study shows that the dynamics of high order structure of chromatin exhibits liquid-like behavior [Maeshima, K., et al., 2016], which could be crucial characteristic in enabling the genome to conduct SOC gene expression control for cell-fate determination.

Further studies on these matters are needed to clarify the underlying fundamental molecular mechanism. The development of a theoretical foundation for the autonomous critical control mechanism in genome expression as revealed in our findings is expected to open new doors for a general control mechanism of the cell-fate determination and genome computing (see Discussion in [Tsuchiya, M, et al, 2015, 2016]), i.e., the existence of ‘*genome intelligence*’.

As for now, we can safely affirm that the strong interactions among genes with very different expression variance and physiological roles push for a complete re-shaping of the current molecular-reductionist view of biological regulation looking for single ‘significantly affected’ genes in the explanation of the regulation processes. The view of the genome acting as an integrated dynamical system is here to stay.

## IV. Methods

### Biological Data Sets

We analyzed mammalian transcriptome experimental data for seven distinct cell fates in different tissues:

#### Cell population

1. Microarray data of the activation of ErbB receptor ligands in human breast cancer MCF-7 cells by EGF and HRG; Gene Expression Omnibus (GEO) ID: GSE13009 (N = 22277 mRNAs; experimental details in [Saeki Y, et al., 2009]) at 18 time points: *t*_1_ = 0, *t*_2_ = 10,15, 20, 30, 45, 60, 90min, 2, 3, 4, 6, 8, 12, 24, 36, 48, *t*_T = 18_ = 72h.
2. Microarray data of the induction of terminal differentiation in human leukemia HL-60 cells by DMSO and atRA; GEO ID: GSE14500 (N = 12625 mRNAs; details in [Huang, S., et al., 2005]) at 13 time points: *t*_1_= 0, *t*_2_ = 2, 4, 8, 12, 18, 24, 48, 72, 96, 120, 144, *t*_T=13_ =168h.

#### Single cell

3. RNA-Seq data of early embryonic development in human and mouse developmental stages in RPKM values; GEO ID: GSE36552 (human: N = 20286 RNAs) and GSE45719 (mouse: N = 22957 RNAs) with experimental details in [Yan, L., et al., 2013] and [Deng, Q., et al., 2014], respectively. We analyzed 7 human and 10 mouse embryonic developmental stages listed below: Human: oocyte (m = 3), zygote (m = 3), 2-cell (m = 6), 4-cell (m = 12), 8-cell (m = 20), morula (m = 16) and blastocyst (m = 30), Mouse: zygote (m = 4), early 2-cell (m = 8), middle 2-cell (m = 12), late 2-cell (m = 10), 4-cell (m = 14), 8-cell (m = 28), morula (m = 50), early blastocyst (m = 43), middle blastocyst (m = 60) and late blastocyst (m = 30), where m is the total number of single cells.
4. RNA-Seq data of T helper 17 (Th17) cell differentiation from mouse naive CD4+ T cells in RPKM (Reads Per Kilobase Mapped) values, where Th17 cells are cultured with anti-IL-4, anti-IFNγ, IL-6 and TGF-β, (details in [Ciofani, M., et al., 2012]; GEO ID: GSE40918 (mouse: N = 22281 RNAs) at 9 time points: *t*_1_ = 0, *t*_2_ = 1, 3, 6, 9, 12, 16, 24, *t*_T=9_ = 48h. For each time point, the reference sample numbers are listed: GSM1004869-SL2653 (t= 0h); GSM1004941-SL1851 (t =1h); GSM1004943-SL1852 (t = 3h); GSM1005002-SL1853 (t= 6h); GSM1005003-SL1854 (t= 9h); GSM1004934-SL1855 (t= 12h); GSM1004935,6,7-SL1856, SL8353, SL8355 (t= 16h; average of three data); GSM1004942-SL1857 (t = 24h); GSM1004960-SL1858 (t= 48h).

In reference to the colors used in the various plots throughout this report, they are based on the experimental events and have been assigned as the following: black as the initial event, purple as the 2^nd^ event, and the subsequent events as blue, dark cyan, dark green, dark yellow, brown, orange, red, dark pink, and pink.

For microarray data, the Robust Multichip Average (RMA) was used to normalize expression data for further background adjustment and to reduce false positives [Bolstad, B. M., et al., 2003; Irizarry, R. A., et al, 2003; McClintick, J. N., Edenberg, H. J., 2006].

For RNA-Seq data, RNAs with RPKM values of zero over all of the cell states were excluded. In the analysis of sandpile criticality, random real numbers in the interval [0, *a*] generated from a uniform distribution were added to all expression values (only in **Figures 5C,6C**). This procedure avoids the divergence of zero values in the logarithm. The robust sandpile-type criticality through the grouping of expression was checked by changing a positive constant: *a* (0 < *a* < 10); we set *a* = 0.001. Note: The addition of large random noise (*a* >> 10) destroys the sandpile CP.

### Normalized Root Mean Square Fluctuation (nrmsf)

*Nrmsf* (see more Methods in [Tsuchiya, M., et al., 2015]) is defined by dividing *rmsf* (root mean square fluctuation) by the maximum of overall {*rmsf*_i_}:

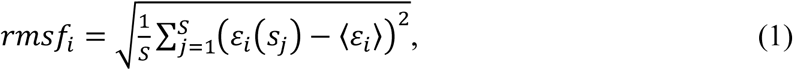

where *rmsf*_*i*_ is the *rmsf* value of the *i*^*th*^ RNA expression, which is expressed as *ε*_*i*_(*s*_*j*_) at a specific cell state *s*_*j*_ or experimental time (e.g., in mouse embryo development, *S* = 10: *s*_1_ *=* zygote, early 2-cell, middle 2-cell, late 2-cell, 4-cell, 8-cell, morula, early blastocyst, middle blastocyst and *s*_10_ *=* late blastocyst), and ⟨*ε*_*i*_⟩ is its expression average over the number of cell states. Note: *nrmsf* is a time-independent variable.

### CM correlation analysis

To investigate the transition dynamics, the correlation metrics based on the center of mass (CM) grouping, the *CM correlation* is built upon the following basic statistical formalization:

1. CM grouping: genome expression is considered as a *N*-dimensional vector, where each expression is subtracted by the average value (CM) of the whole expression at *t* = *t*_j_ (refer to **Figure 1**). Next, the whole expression is sorted and grouped according to the degree of *nrmsf*, where CM grouping has *K* groups and within each group there are *n* number of expressions (*N* = *K*.*n*): *N*-dimensional CM grouping vector, ***C***(*t*_*j*_) = (***c***^1^(*t*_*j*_), ***c***^2^(*t*_*j*_), …, ***c***^*k*^(*t*_*j*_),.., ***c***^*K*^(*t*_*j*_)) ; ***c***^1^(*t*_*j*_) and ***c***^*K*^(*t*_*j*_) are the highest and lowest group vectors of *nrmsf*, respectively. Here, the unit vector of *k*^th^ vector ***c***^*k*^(*t*_*j*_) is defined as 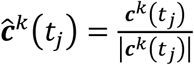. Note that the less than *n* elements in the last group (the lowest *nrmsf*) have been removed from the analysis.
2. Keeping in mind, correlation corresponds to the cosine of angle between unit vectors, i.e., inner product of unit vectors (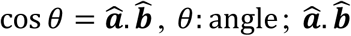: dot product (scalar) of unit 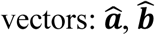).

Two different CM correlations can be considered:

i. **Spatial CM correlation**: for a given time point (*t* = *t*_*j*_), development of CM correlation between the first group (highest *nrmsf* group) and other vectors: ***ĉ***^1^(*t*_*j*_). ***ĉ***^*k*^(*t*_*j*_); (*k* = 2,3,..*K*).
ii. **Temporal CM correlation**: for a given group (*k*), development of CM correlation between the initial and other experimental points: ***ĉ***^*k*^(*t*_1_). ***ĉ***^*k*^(*t*_*j*_) (*k*= 1,2,3,..*K*) over experimental time points, *t*_*j*_ (see **Biological Data Sets**).

### *Probability density function* (PDF)

By means of density analysis of noisy gene-expression profiles, the robustness of gene expression clustering was demonstrated [Shu, G., at al., 2003]. The probability density function (PDF) based on Gaussian kernel is examined. We consider *N*-dimensional whole expression vector for natural logarithm (natural-log) of expression and its fold-change vector, where natural-log of whole expression vector is defined as *ln*(***ε***(*t*_j_)) = (*ln*(*ε*_1_(*t*_j_)), *ln*(*ε*_2_(*t*_j_)),…, *ln*(*ε*_*N*_(*t*_j_))) with the *i*^th^ expression, *ε*_*i*_(*t*_j_) at *t* = *t*_j_ (**Figure 1**) and its fold-change vector is defined as *ln*(***ε***(*t*_*j*_)/***ε***(*t*_*k*_)) = (*ln*(*ε*_1_(*t*_*j*_)/*ε*_1_(*t*_*k*_)), *ln*(*ε*_2_(*t*_*j*_)/*ε*_2_(*t*_*k*_)),…, *ln*(*ε*_*N*_(*t*_*j*_)/*ε*_*N*_(*t*_*k*_)). This stratifies the logarithm of a quotient: *ln*(***ε***(*t*_*j*_)/***ε***(*t*_*k*_)) = *ln*(***ε***(*t*_j_)) - *ln*(***ε***(*t*_*k*_)). Next, we consider these vectors from their CM, CM natural-log of whole expression vector and it fold-change vector. CM natural-log of whole expression vector is defined as *ln*(***c***(*t*_j_)) = *ln*(***ε***(*t*_j_)) - CM(*ln*(***ε***(*t*_j_))) and 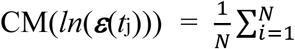 and its fold-change vector as *ln*(***c***(*t*_j_)/***c***(*t*_*k*_)) = *ln*(***ε***(*t*_*j*_)/***ε***(*t*_*k*_)) - CM(*ln*(***ε***(*t*_*j*_)/***ε***(*t*_*k*_))). This vector representation again satisfies the logarithm of a quotient: *ln*(***c***(*t*_*j*_)/***c***(*t*_*k*_)) = *ln*(***c***(*t*_j_)) - *ln*(***c***(*t*_*k*_)).

From embryo to cancer development, global avalanches through the change in the CP are captured and furthermore, scaling behaviors and travelling expression wave on scaling behavior in human and mouse embryo development (RNA-Seq data) is revealed (see **Figures 7C, 8C**), which suggests coordinated DNA folding-unfolding dynamics.

### Expression Flux Analysis

We developed the expression flux analysis to describe genome engine mechanism on both single cell and population genome expression. The key fact is that the dynamics of coherent behavior emerged from stochastic expression in distinct critical states (coherent-stochastic behavior: CSB) follows the dynamics of the CM of a critical state. This convergence shows that the CM of a critical state acts as a critical-state attractor for stochastic expression within the critical state. We have developed the expression flux approach to reveal dynamic interaction flux between critical-state attractors [Tsuchiya, M., et al., 2016-2018].

The CSB in a critical state corresponds to the scalar dynamics of its CM. The numerical value of a specific critical state (i.e., super-, near- or sub-critical state) is represented by *X*(*s*_j_) at a specific experimental event (*s*_j_), where an experimental event (*s*_*j*_) corresponds to a cell state or an experimental time point. The expression flux between critical states is interpreted as a non-equilibrium system and evaluated in terms of a dynamic network of effective forces, where interaction flux is driven by effective forces between different critical states and can be described by a second-order time difference. It is important to note that the oscillatory phenomenon is interpreted using a second-order difference equation with a single variable. This is equivalent to inhibitor-activator dynamics given by a couple of first-order difference equations with two variables. Flux dynamics approach is further developed to analyze quantitative evaluation of the degree of non-harmonicity and time reversal symmetry breaking in nonlinear coupled oscillator systems [Tsuchiya, M., et al., 2018].

Basic formulas of expression flux dynamics are given as follows:

#### Net self-flux of a critical state

The net self-flux, the difference between the IN flux and OUT flux, describes the effective force on a critical state. This net self-flux represents the difference between the positive sign for incoming force (net IN self-flux) and the negative sign for outgoing force (net OUT self-flux); the CM from its average over all cell states represents up- (down-) regulated expression for the corresponding net IN (OUT) flux.

The effective force is a combination of incoming flux from the past to the present and outgoing flux from the present to the future cell state:

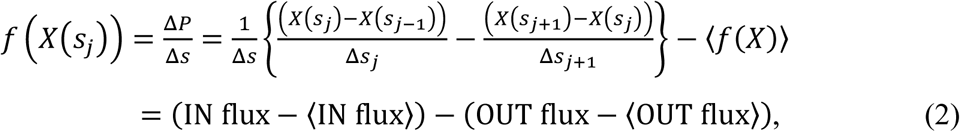

where Δ*P* is the change in momentum with a unit mass (i.e., the impulse: *F*Δ*s* = Δ*P*) and natural log of average (<…>) of a critical state, 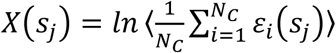 with the *i*^th^ expression *ε*_*i*_(*s*_*j*_) at the *j*^th^ experimental event, *s* = *s*_j_ (*N*_*C*_ = the number of RNAs in a critical state; refer to **Tables 1,2**); the average of net self-flux over the number of critical states, <*f*(X)> = <INflux> - <OUTflux>.

Here, scaling and critical behaviors occur in log-log plots of group expression, where the natural log of an average value associated with group expression such as *ln*<*nrmsf*> and *ln*<*expression*> is taken. Thus, in defining expression flux, the natural log of average expression (CM) of a critical state is considered.

It is important to note that each embryo cell state is considered as a statistical event (note: a statistical event does not necessarily coincide with a biological event) and its development as time arrow (time-development) when evaluating the average of group expression: fold change in expression and temporal expression variance (*nrmsf*). This implies that an interval in the dynamical system (Equation (2)) is evaluated as difference in event, i.e., Δ*s*_j_ = *s*_j_ - *s*_j-1_= 1 and Δ*s* = *s*_j+1_ - *s*_j-1_ *=* 2 in embryo development, as well as difference in experimental times such as in cell differentiation (note: actual time difference can be considered as scaling in time). Then, we evaluate a force-like action in expression flux.

#### The interaction flux of a critical state

The interaction flux represents flux of a critical state *X*(*s*_j_) with respect to another critical state (Super, Near, Sub) or the environment (E: milieu) *Y*_*j*_ can be defined as:

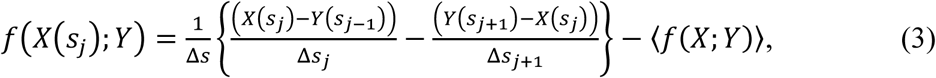

where, again, the first and second terms represent IN flux and OUT flux, respectively, and the net value (i.e., IN flux-OUT flux), represents incoming (IN) interaction flux from *Y* for a positive sign and outgoing (OUT) interaction flux to *Y* for a negative sign. *Y* represents either the numerical value of a specific critical state or the environment, where a state represented by *Y* is deferent from one by *X.*

With regard to the global perturbation event, the net **kinetic energy flux** [Tsuchiya, M., et al., 2016] clearly reveals it in a critical state (**Figure 11**):

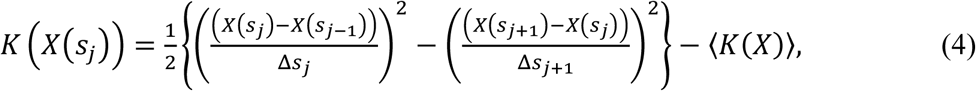

where the kinetic energy of the CM for the critical state with unit mass at *s* = *s*_*j*_ is defined as 1/2. *v*(*s*_*j*_)_2_ with average velocity: 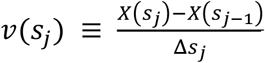.

#### Net self-flux as summation of interaction fluxes

Due to the law of force, the net self-flux of a critical state is the sum of the interaction fluxes with other critical states and the environment:

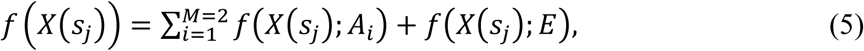

where state *A*_*i*_ ∈ {Super, Near, Sub} with *A*_*i*_ ≠ *X*, and *M* is the number of internal interactions (*M* = 2), i.e., for a given critical state, there are two internal interactions with other critical states. Equation (5) tells us that the sign of the difference between the net self-flux and the overall contribution from internal critical states, 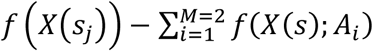, reveals incoming flux (positive) from the environment to a critical state or outgoing flux (negative) from a critical state to the environment; when the difference in all critical states becomes zero, the genome system itself lies in a stable point (*non-equilibrium fixed point*) of thermodynamically open system (no average flux flow from the environment). Average between-state flux (**Figure 11**) represents a stable manifold of the thermodynamically open system.

Here, we need to address the previous result of expression flux dynamics in mouse single-cell genome expression [Tsuchiya, M., et al., 2017], where expression of a critical state was taken as 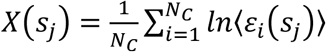, which has different ordering of operations: first taking the natural log of expression and then, average operation. Hence, in flux dynamics, we examine whether or not mathematical operation between averaging and natural log, i.e., operation between *In* 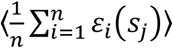 and 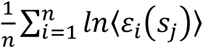 can be exchanged (mathematically commuted). In microarray data, flux behaviors do not change much between these action ordering (almost the same: commuted). Whereas in RNA-Seq data, they are not commuted due to its data structure with lots of zero values; adding small random noise into log of expression, *ln*⟨*ε*_*i*_(*s*_*j*_)⟩ (previous result) makes good effect (noise-sensitive), but not in *In* 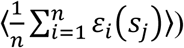: noise-insensitive (this report). Although detail dynamics of interaction flux changes by taking different action-orderings in RNA-Seq data (e.g., Fig. 6 in [Tsuchiya, M., et al., 2017]), two important characteristics in genome-engine: the formation of dominant cyclic flux between super- and sub-critical states and the generator role of the sub-critical state do not change (invariant features). Thus, we conclude that the concept of the genome-engine is quite robust.

## Contributions

MT initiated the project; MT, AG and KY designed the study; MT developed the study and analyzed data; AG and KY provided theoretical support; MT, AG and KY wrote the manuscript.

## Acknowledgments

This manuscript has been released as a Pre-Print [Tsuchiya, M., Giuliani, A., Yoshikawa, K., 2020]. MT sincerely thanks the following institution and individuals who helped complete this research project: the SEIKO Life Science Laboratory, Osaka, Japan, his family (particularly, his daughters, Drs. Kimiko and Kazumi Tsuchiya, and Dr. Harry Taylor with any editing), Drs. Andrzej Kasperski for fruitful discussions through the proof-reading and Jekaterina Erenpreisa for biological discussions.

---

**Supplementary Figure S1.**
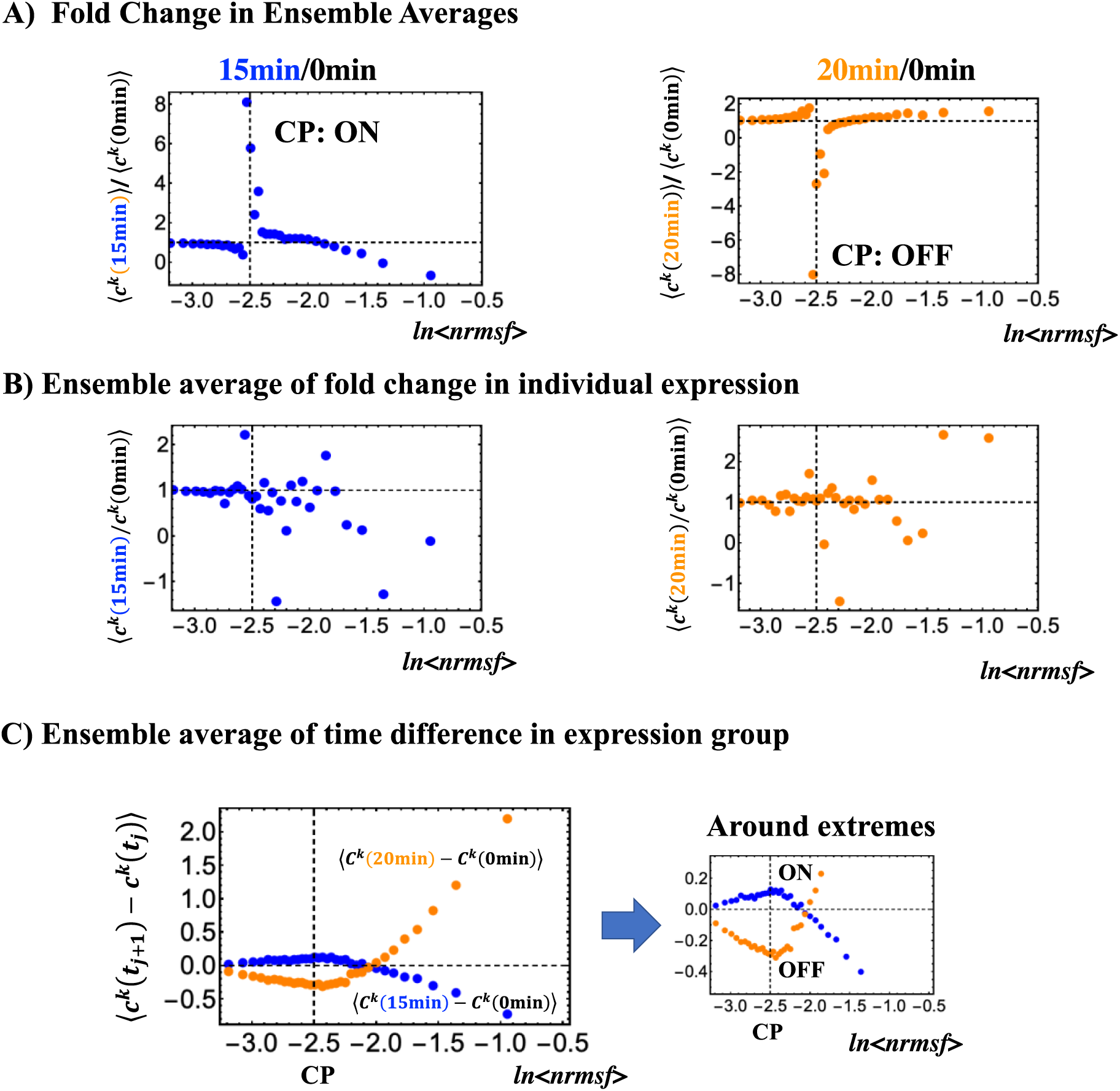
(HRG-stimulated MCF-7 cells): **A)** Fold change in ensemble (group) averages between expression groups shows that coherent-stochastic behavior (emergent coherent behavior from stochastic expression) represented by the center of mass (average) of group reveals a clear coil-globule transition. **B)** Ensemble average of fold change in individual expressions between two temporal groups, <***c***^*k*^(t_j+1_)/***c***^*k*^(t_j_)>, does not reveal any transitional behavior that is attributable to the stochastic behavior of expression (sensitive in fold change). **C)** Ensemble average of time difference in expression groups, <***c***^*k*^(t_j+1_)- ***c***^*k*^(t_j_)>, supports the coherent scenario in (**A**), where at around the CP (*ln*<*nrmsf*>∼ -2.5) there is a positive maximum at 0-15min and negative minimum at 0-20min that corresponds to either an ON or OFF state, respectively.

## References

1. Bak, P., Tang, C. Wiesenfeld, K. (1987). Self-organized criticality: An explanation of the 1/f noise. Phys. Rev. Lett. 59: 381–384; doi: 10.1103/PhysRevLett.59.381.

2. Bolstad, B. M., Irizarry, R. A., Astrand, M., Speed, T. P. (2003). A comparison of normalization methods for high density oligonucleotide array data based on variance and bias. Bioinformatics 19, 185–193; doi: 10.1093/bioinformatics/19.2.185.

3. Censi, F., Giuliani, A., Bartolini, P., Calcagnini, G. (2011). A Multiscale Graph Theoretical Approach to Gene Regulation Networks: A Case Study in Atrial Fibrillation. IEEE Trans. Biomed. Eng. 99, 1–5; doi: 10.1109/TBME.2011.2150747.

4. Ciofani, M., Madar, A., Galan, C., Sellars, M., Mace, K., Pauli, F., Agarwal, A., Huang, W., Parkhurst, C. N., Muratet, M., et al. (2012). A validated regulatory network for Th17 cell specification. Cell 151, 289–303; doi: 10.1016/j.cell.2012.09.016.

5. Deng, Q., Ramsköld, D., Reinius, B., Sandberg, R. (2014). Single-cell RNA-seq reveals dynamic, random monoallelic gene expression in mammalian cells. Science 343, 193–196; doi: 10.1126/science.1245316.

6. Flory, P. (1953). Principles of Polymer Chemistry, Cornell University Press.

7. Gennes, P.G. (1979). Scaling Concepts in Polymer Physics, Cornell University Press, Ithaca, NY.

8. Giuliani, A., Tsuchiya, M., Yoshikawa, K. (2018). Self-Organization of Genome Expression from Embryo to Terminal Cell Fate: Single-Cell Statistical Mechanics of Biological Regulation. Entropy 20, 13; https://doi.org/10.3390/e20010013.

9. Halley, J. D., Burden, F. R., Winkler, D. A. (2009). Summary of stem cell decision making and critical-like exploratory networks. Stem Cell Res. 2, 165–177; https://doi.org/10.1016/j.scr.2009.03.001.

10. Huang, S., Eichier, G., Bar-Yam, Y., Ingber, D. E. (2005), Cell fates as high-dimensional attractor states of a complex gene regulatory network. Phys. Rev. Lett. 94, 128701–128705; doi: 10.1103/PhysRevLett.94.128701.

11. Irizarry, R. A., Hobbs, B., Collin, F., Beazer-Barclay, Y. D., Antonellis, K. J., Scherf, U., Speed, T. P. (2003). Exploration, normalization, and summaries of high density oligonucleotide array probe level data. Biostatistics 4, 249–264; DOI: 10.1093/biostatistics/4.2.249

12. Jensen, H. J. (1998). Self-Organized Criticality. Cambridge Univ. Press, UK.

13. Kauffman, S. A. (1993). The Origins of order self-organization and selection in evolution. Oxford Univ. Press, New York; https://doi.org/10.1142/9789814415743_0003.

14. Kim, K.Y., Wang, J. (2007). Potential energy landscape and robustness of a gene regulatory network: Toggle switch. PLoS Comp. Biol. 3, e60; https://doi.org/10.1371/journal.pcbi.0030060.

15. Langton, C. G. (1990). Computation at the edge of chaos - phase transitions and emergent computation. Physica D 42, 12–37; https://doi.org/10.1016/0167-2789(90)90064-V.

16. MacArthur, B. D., Ma’ayan, A., Lemischka, I. R. (2009). Systems biology of stem cell fate and cellular reprogramming. Nat. Rev. Mol. Cell Biol. 10, 672–681; doi: 10.1038/nrm2766.

17. Maeshima, K., Ide, S., Hibino, K., Sasai, M. (2016). Liquid-like behavior of chromatin. Curr. Opi. Gene. Dev. 37, 36–45; https://doi.org/10.1016/j.gde.2015.11.006.

18. Marković, D., Gros, C. (2014). Power laws and self-organized criticality in theory and nature. Phys. Rep. 536, 41–74; https://doi.org/10.1016/j.physrep.2013.11.002.

19. McClintick, J. N., Edenberg, H. J. (2006). Effects of filtering by present call on analysis of microarray experiments. BMC Bioinformatics 7, 49; doi: 10.1186/1471-2105-7-49.

20. Mojtahedi, M., Skupin, A., Zhou, J., Ivan Castaño, I. G., Leong-Quong, R. Y., Chang, H., Giuliani, A., Huang, S. (2016). Cell Fate Decision as High-Dimensional Critical State Transition. PLoS Biol 14: e2000640; doi: 10.1371/journal.pbio.2000640.

21. Muñoz, M. A. (2018). Colloquium: Criticality and dynamical scaling in living systems. Rev. Mod. Phys. 90, 031001–031030; https://doi.org/10.1103/RevModPhys.90.031001.

22. Nagashima T, Shimodaira H, Ide K, Nakakuki T, Tani Y, Takahashi K, et al. (2007) Quantitative transcriptional control of ErbB receptor signaling undergoes graded to biphasic response for cell differentiation. J. Biol. Chem. 282: 4045–4056; doi: 10.1074/jbc.M608653200.

23. Nakai, T., Hizume, K., Yoshimura, S. H., Takeyasu, K., Yoshikawa, K. (2005). Phase transition in reconstituted chromatin *Europhys*. Lett. 69, 1024–1030; https://doi.org/10.1209/epl/i2004-10444-6.

24. Nakakuki T, Birtwistle MR, Saeki Y, Yumoto N, Ide K, Nagashima T, et al. (2010) Ligand-specific c-Fos expression emerges from the spatiotemporal control of ErbB network dynamics. Cell 141: 884–896; doi: 10.1016/j.cell.2010.03.054.

25. Raser, J. M., O’Shea, E. K. (2005). Noise in gene expression: Origins, consequences, and control. Science 309, 2010–2013; doi: 10.1126/science.1105891.

26. Saeki, Y., Endo, T., Ide, K., Nagashima, T, Yumoto, N., Toyoda, T., Suzuki, H., Hayashizaki, Y., Sakaki, Y., Okada-Hatakeyama M. (2009). Ligand-specific sequential regulation of transcription factors for differentiation of MCF-7 cells. BMC Genomics 10: 545; doi: 10.1186/1471-2164-10-545.

27. Sakaue, T., Yoshikawa, K. (2006). On the formation of rings-on-a-string conformations in a single polyelectrolyte chain: A possible scenario. J. Chem. Phys. 125, 32767–6; doi: 10.1063/1.2244555.

28. Schiessel, H. (2003). The physics of chromatin. J. Phys. Condes. Matter, 15, R699–R774; doi: 10.1088/0953-8984/27/6/060301.

29. Shew, C.Y., Yoshikawa, K. (2007). Mean field theory for the intermolecular and intramolecular conformational transitions of a single flexible polyelectrolyte chain. J. Chem. Phys. 126, 32767–9; DOI: 10.1063/1.2714552.

30. Shu G, Zeng B, Chen YP, Smith OH (2003) Performance assessment of kernel density clustering for gene expression profile data. Comp Funct Genomics 4, 287–299; https://doi.org/10.1002/cfg.290

31. Suzuki, Y., Yoshikawa, Y., Yoshimura, S.H., Yoshikawa, K., Takeyasu, K. (2011). Unraveling DNA dynamics using atomic force microscopy. Wiley Interdiscip. Rev. Nanomed. Nanobiotechnol., 3, 574–588; doi: 10.1002/wnan.150.

32. Takahashi, K., Yamanaka, S. (2016). A decade of transcription factor-mediated reprogramming to pluripotency. Nature Rev. Mol. Cell Biol. 17, 183–193; doi: 10.1038/nrm.2016.8.

33. Tsuchiya, M., Wong, ST., Yeo, ZX., Colosimo, A., Palumbo, MC., Crescenzi, M., Mazzola, A., Negri, R., Bianchi, MM., Selvarajoo, K., Tomita, M., Giuliani, A. (2007). Gene expression waves: cell cycle independent collective dynamics in cultured cells. FEBS J. 274, 2874–2886; doi: 10.1111/j.1742-4658.2007.05822.x.

34. Tsuchiya, M., Hashimoto, M., Takenaka, Y., Motoike, I. N., Yoshikawa, K. (2014). Global genetic response in a cancer cell: Self-organized coherent expression dynamics. PLOS One 9: e97411; https://doi.org/10.1371/journal.pone.0105491.

35. Tsuchiya, M., Giuliani, A., Hashimoto, M., Erenpreisa, J., Yoshikawa, K. (2015). Emergent Self-Organized Criticality in gene expression dynamics: Temporal development of global phase transition revealed in a cancer cell line. PLoS One 11, e0128565; https://doi.org/10.1371/journal.pone.0128565.

36. Tsuchiya, M., Giuliani, A., Hashimoto, M., Erenpreisa, J., Yoshikawa, K. (2016). Self-organizing global gene expression regulated through criticality: Mechanism of the cell-fate change. PLoS ONE 11: e0167912; https://doi.org/10.1371/journal.pone.0167912.

37. Tsuchiya, M., Giuliani, A., Yoshikawa, K. (2017). Single-Cell Reprogramming in Mouse Embryo Development through a Critical Transition State. Entropy 19, 584; https://doi.org/10.3390/e19110584.

38. Tsuchiya, M., Giuliani, A., Yoshikawa, K. (2018). A Quantitative Evaluation of Symmetry Breaking In Nonlinear-Oscillatory System - Based on Flux Dynamics (Effective force) View Point. Presentation; doi: 10.13140/RG.2.2.34048.74240.

39. Tsuchiya, M., Giuliani, A., Yoshikawa, K. (2020). Cell-Fate Determination from Embryo to Cancer Development: Genomic Mechanism Elucidated. Preprint, bioRxiv.

40. Yan, L., Yang, M., Guo, H., Yang, L., Wu, J., Li, R., Liu, P., Lian, Y., Zheng, X., Yan, J., et al. (2013). Single-cell RNA-Seq profiling of human preimplantation embryos and embryonic stem cells. Nat. Struct. Mol. Biol. 20, 1131–1139; doi: 10.1038/nsmb.2660.

41. Yoshikawa, K. (2002). Field hypothesis on the self-regulation of gene expression. J. Biol. Phys. 28: 701–712; doi: 10.1023/A:1021251125101.

42. Yoshikawa, K., Takahashi, M., Vasilevskaya, V.V., Khokhlov, A.R. (1996). Large Discrete Transition in a Single DNA Molecule Appears Continuous in the Ensemble. Phys. Rev. Lett., 76, 3029-3031; 10.1103/PhysRevLett.76.3029.

43. Yoshikawa, K., Yoshikawa, Y. (2002). Compaction and Condensation of DNA, in Pharmaceutical Perspective of Nucleic Acid-Base Therapy. Tayler & Francis, Abingdon, UK, 137–163.

44. Wagner, J. R., Lee, C. T., Durrant, J. D., Malmstrom, R. D., Feher, V. A., Amaro, R. E. (2016). Emerging Computational Methods for the Rational Discovery of Allosteric Drugs Chem. Rev., 116, 6370–90; doi: 10.1021/acs.chemrev.5b00631.

45. Zimatore, G., Tsuchiya, M., Hashimoto, M., Kasperski, A., Giuliani, A. (2019). The existence of a potential general mechanism of cell-fate change shared by different biological processes. Preprint. bioRxiv; doi: https://doi.org/10.1101/852681.

46. Zinchenko, A., Pyshkina, O., Lezov, A., Sergeyev, V., Yoshikawa, K. (2008). Single DNA molecules: compaction and decompaction (chapter 3), in DNA interactions with polymers and surfactants. Wiley-Blackwell.

